# Multimodal characterization of putative *pan*-inhibitory viral constructs in macaque monkeys

**DOI:** 10.64898/2026.10.27.761178

**Authors:** M. Medalla, P. O. Boucher, C. A. Mojica, H. Bhatt, B. Snyder, Y. Zhou, N. Sengupta, A. I. More, M. Yahaya, X. Opitz-Araya, T. Wang, EK. Lee, D. L. Rosene, R. J. Rushmore, T. L. Moore, B.V. Zemelman, J. T. Ting, B. P. Levi, J. I. Luebke, C. Chandrasekaran

## Abstract

Transgenic approaches and validated viral vector toolboxes have advanced our understanding of how inhibitory neurons shape cognition and motor control in mice and other rodents. However, such advances have been elusive in non-human primates (NHPs). While many studies demonstrate expression of reporter proteins, few studies have provided clear quantitative evidence of whether these viral constructs can be reliably used for assessment of *in vivo* and *in vitro* function in primates, which, in turn, has hindered widespread adoption by NHP researchers. Here, we characterize putative pan-inhibitory cell targeting vectors engineered with regions from the *Dlx5-6* intergenic domain. Pan-inhibitory viral vectors are attractive because they enable ground-truth optotagging and modulation of multiple inhibitory subtypes in the same animal and, when combined with machine learning approaches, enable cell-type classifiers for NHPs. We combined *in vivo* anesthetized electrophysiological recordings, *in vitro* slice electrophysiology, and immunohistochemistry across seven rhesus macaque monkeys to assess the specificity, completeness and efficacy of three constructs in multiple sensory and higher-order association cortices. The vectors reliably restricted transgene expression to inhibitory neurons (81-97%). However, of the local GABAergic neurons only 32-64% expressed the transgene, depending on the construct. This incompleteness arose in part from uneven coverage across subtypes, with strong transgene expression in parvalbumin-positive interneurons (∼60-90%), sparse expression in calbindin-positive interneurons (∼1-7%), and near-complete avoidance of calretinin-positive interneurons. A construct pairing DLX2.0 with ChrimsonR was excellent for *in vivo* optotagging, while h56D paired with bReaChES most effectively suppressed neural activity *in vivo*. In the slice, activating terminals of transgene-expressing neurons optically evoked inhibitory currents in coupled pyramidal neurons confirming transsynaptic functionality of these constructs and their use for inhibitory circuit-specific synaptic assessments and manipulation. Collectively, the results provide rigorous functional characterization, and quantitative guidance for NHP researchers seeking to deploy GABAergic cell-type-specific tools in their experiments.

## 1. Introduction

Inhibitory neurons are fundamental components of brain circuits involved in cognition and motor control^1,2–4^. Studies in mice often leverage transgenic approaches and/or validated viral toolboxes to optogenetically identify (i.e., ‘optotag’) inhibitory neurons or to modulate their activity^3,4^. Such efforts have enabled rapid progress in delineating circuit motifs and the role of various inhibitory neuron types in rodent behavior^3,4,5,6^. In contrast, there has been comparatively limited progress in understanding the role of inhibitory neurons in behaving non-human primates (NHPs)^7^, despite reports of promising constructs driven by GABAergic neuron-specific enhancers^8–15^.

Several factors contribute to this lack of progress. Studies are often proof of concept and primarily aim to demonstrate that enhancers can selectively drive reporter protein expression in a small number of NHPs (typically 1-2) and brain regions^8–10^. Inclusion of each additional component that a cell must express (e.g., opsin, reporter) from a viral construct increases the probability for toxic effects or could change the function of the cell^16,17^. Thus, whether neurons remain viable (i.e., do they survive), functional (i.e., do they inhibit), and identifiable (i.e., do they maintain similar spontaneous and optically-evoked waveforms) post-infection with a viral construct that expresses in inhibitory neurons in a NHP brain remains unknown. Moreover, there is only a modest amount of data on the efficacy of these viral constructs to optotag or modulate neurons *in vivo*^7,18^. A further concern is that enhancers are often identified from mouse genomes. Given evolutionary divergence between species, there is no guarantee that a mouse-derived enhancer will perform reliably in NHP species^12^. The cost, effort, and experimental risk in deploying uncharacterized constructs in well-trained NHPs that routinely perform sophisticated tasks is considerable. Without robust quantitative data on real-world performance and detailed qualitative insights on how viral constructs behave *in vivo*, NHP researchers worldwide are understandably reluctant to readily adopt such tools^16,19^.

Viral constructs with pan-inhibitory enhancers would ameliorate some of these issues and accelerate the study of inhibitory neurons. Such constructs would enable optotagging of multiple inhibitory neuron subtypes in the same animal, and area-wide suppression of excitatory neuron activity^20^. In addition, these tools would enable tests of hypotheses of how excitatory and inhibitory neurons combine to drive computation^21^, perception^4,12,15^, cognition^22,23^ and motor control^6^. Combined with machine learning approaches capable of clustering waveforms and spiking patterns by cell type^24–27^, such a construct could further enable subtype separation without requiring subtype-specific tools.

Motivated by this potential, several groups have focused on enhancers of the distal-less homeobox (*Dlx*) gene family. These enhancers are conserved regulatory elements located between the paired genes *Dlx5* and *Dlx6* which encode transcription factors essential for GABAergic interneuronal differentiation and migration in the developing forebrain (Fig. S1a)^28–31^. The first studies adapted these regions of the mouse genomes to build putative pan-inhibitory enhancers for use across species (e.g., mDlx, or closely related hDlx), which surprisingly labeled interneurons in adult animals despite the postnatal decline in Dlx5/6 expression^32^. Other studies boosted expression by repeating 3 core elements of the conserved portions of the hDlx enhancer in tandem to improve expression in GABAergic neurons (Fig. S1b)^10^. Yet another study identified a different human sequence based on mouse-human conservation alone (i.e., h56D^9^) to develop other pan-inhibitory enhancers (Fig. S1c).

Evidence on how these constructs actually perform in primates is promising but uneven. In marmosets, reporter-only versions of Dlx-based enhancers label inhibitory neurons in the visual cortex with high specificity (of all virus-labeled cells, what fraction are the intended cell type) and reasonable completeness ^8,9,13^ (of all cells belonging to an intended target population, what fraction did the virus label). In macaques, one study demonstrated high specificity of Dlx-based enhancers for GABAergic neurons in ex-vivo tissue culture, and others have used these enhancers to modulate and record from inhibitory neurons in visual^7,18^ and motor cortices^33^. *in vitro* functional validation at the synaptic level is largely unavailable in primates. Finally, studies in the developing mouse brain^28–31^ and recent work in macaque V1^11^ raises the possibility that Dlx enhancers favor GABAergic neurons arising from particular progenitor pools rather than the inhibitory population as a whole^28–31^. Thus, whether these pan-inhibitory viral tools label, optotag, and functionally engage inhibitory neurons across the heterogeneous cytoarchitecture of the macaque cortex therefore remains open.

The current study addresses this open question with a novel multimodal approach. We characterize the specificity, completeness, and, crucially, the *in vivo* and *in vitro* functional efficacy of three recently developed constructs that combine Dlx-based enhancers with opsin-reporter fusion proteins to restrict transgene expression to inhibitory neurons^9,10^. We combine *in vivo* anesthetized electrophysiological recordings to optotag inhibitory neurons, *in vitro* slice electrophysiology to stimulate opsin-expressing inhibitory neurons and assess suppression of coupled excitatory neurons, and immunohistochemistry to assess cell health and the inhibitory subtypes expressing the transgene. We localized our injections to four functionally and cytoarchitectonically distinct cortical areas: primary sensory (primary somatosensory, S1), premotor (dorsal premotor, PMd), prefrontal (lateral prefrontal cortex, LPFC, a46), and limbic (mid-cingulate cortex, MCC, a24c/d).

We found that these constructs reliably restricted transgene expression to inhibitory neurons. However the viral constructs preferentially expressed opsins and reporters in parvalbumin-positive (PV+) and some calbindin-positive (CB+, e.g., somatostatin-positive (SST+)) neurons, but never in calretinin-positive (e.g., vasoactive intestinal peptide-positive (VIP+)) neurons. This pattern is consistent with a recent finding in rat and macaque cortex demonstrating that *Dlx* enhancer-based vectors are more selective for PV-expressing subtypes^11^, but contrasts with their pan-inhibitory expression in mice^8–10^. Further, we demonstrated that these constructs successfully optotag inhibitory neurons *and* suppress the activity of putative excitatory neurons *in vivo* and pyramidal neurons *in vitro*, confirming intact inhibitory functionality. Collectively, these results provide a rigorous functional characterization of putative pan-inhibitory viral constructs in the macaque and offer quantitative insight for NHP researchers seeking to deploy such viral tools in their experiments.

## 2. Results

### 2.1. Multi-modal approach to characterize performance of pan-inhibitory viral constructs

At a high level, we developed a first-of-its-kind multimodal approach to validate and test various pan-inhibitory viral constructs for optically modulating inhibitory neurons *in vivo* and *in vitro* in the monkey (Fig. 1a). First monkeys underwent a survival surgery where we performed a craniotomy and reflected the dura. We then identified relevant brain areas using visual landmarks and injected relevant viral constructs (Fig. S2a, S4). After an average incubation period of ∼13 weeks (Fig. S3), the animals underwent a procedure where we optically stimulated the opsin-expressing neurons *in vivo* and recorded the responses to identify optotagged units (Fig. S2b). We used Kilosort4^34^ to spike sort all recorded units and then identified the units whose activity increased or decreased in response to the optical stimulation (Fig. S2c). At the end of the procedure we euthanized and perfused the animals to obtain various blocks of tissue for *in vitro* slice electrophysiology (Fig. S2d,e) and subsequent immunohistochemistry (Fig. S2f).

**Figure 1:**
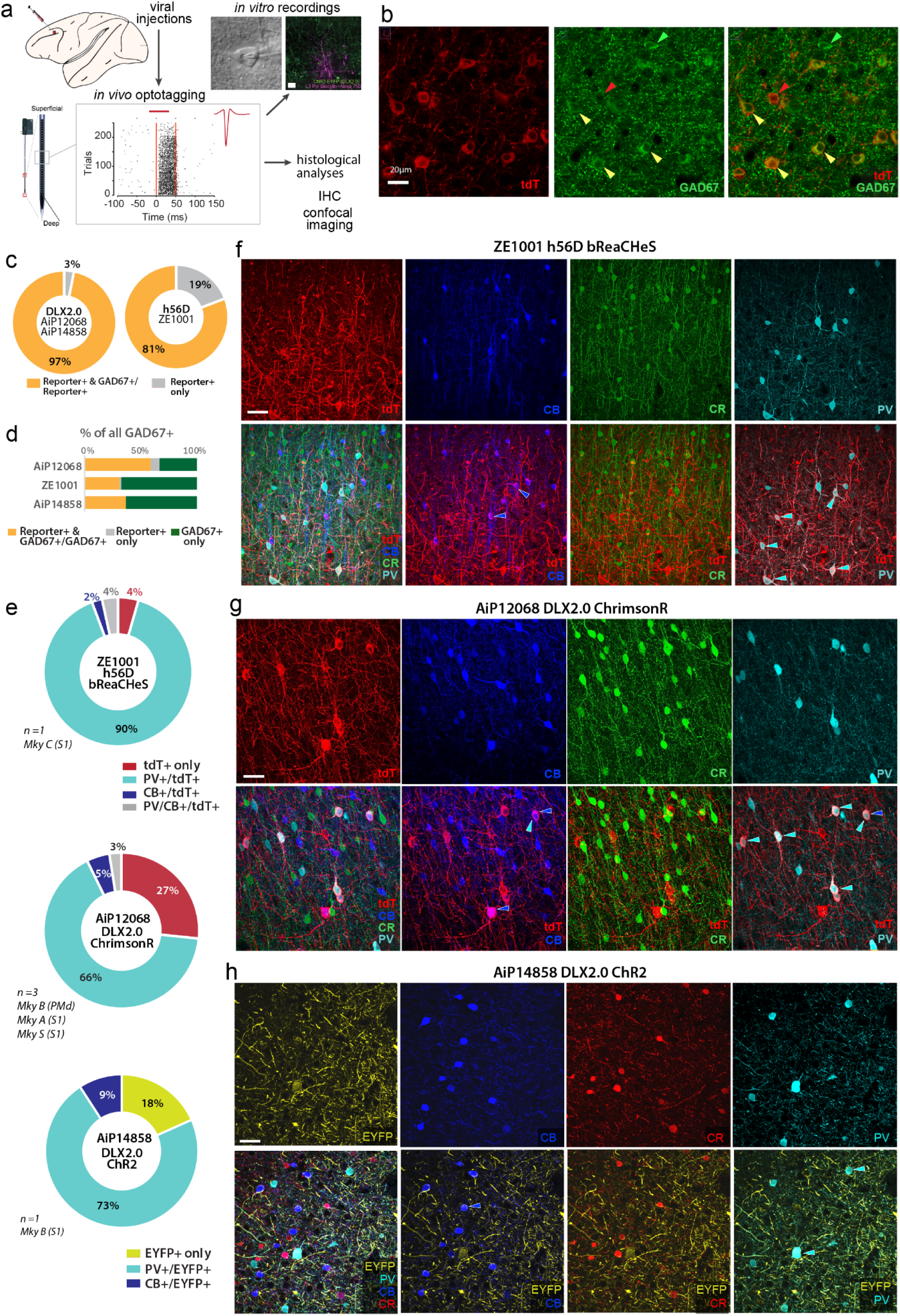
All ‘pan’-inhibitory constructs demonstrate a selective bias for PV+ neurons. (**a**) Multimodal validation of 3 viral constructs to target and functionally control GABAergic neurons. We surgically injected viral constructs into cortical brain areas (e.g., PMd). After a ∼13 week survival, we performed *in vivo* anesthetized recordings to identify optotagged units. We harvested brain tissue from the left hemisphere areas containing the viral injections for *in vitro* recordings, and we fixed the remaining intact right hemisphere for injection site validation and histological analyses of transgene-expressing neurons. (**b**) Representative 40x Z-max projection of merged and individual channel fluorescent images showing co-labeling of tdTomato (tdT, red), construct reporter (AiP12068), with Glutamic Acid Decarboxylase 67 (GAD67, green), a marker for GABAergic cells. Red arrowheads indicate tdT+ cells, green arrowheads indicate GAD67+ cells and white arrowheads indicate tdT+/GAD67+ cells (i.e., transgene-expressing GABAergic cells). (**c**) Ring charts of reporter+ and GAD67+ coexpression for DLX2.0 (AiP12068 and AiP14858) and h56D bReaChES (ZE1001) constructs. DLX2.0 shows high specificity for GAD67+ cells, while h56D bReaChES shows broader expression including GAD67-cells. (**d**) Stacked bar charts showing the transgene-expressing reporter+ neurons as a proportion of the total GAD67+ GABAergic population (reporter+ & GAD67+/GAD67+, reporter+ only, and GAD67+ only) for each construct. (**e**) Ring charts of subtype specificity (percent of reporter+ neurons co-expressing CB, PV, or CR) for ZE1001 h56D-bReaChES-tdT, AiP12068 DLX2.0-ChrimsonR-tdT, and AiP14858 DLX2.0-ChR2-EYFP constructs respectively. ZE1001 shows predominant labeling of PV+ interneurons with smaller contributions from CB+ and mixed populations (top). AiP12068 shows majority expression in PV+ interneurons with additional labeling of tdT-only and minor CB+ populations (middle). AiP14858 shows predominant targeting of PV+ interneurons with smaller fractions of EYFP-only and CB+ populations (bottom). (**f-h**) Representative multi-channel confocal fluorescent image stacks of reporter proteins (red tdT, yellow EYFP) together with interneuron subtype markers: calbindin (CB, blue), calretinin (CR, green), and parvalbumin (PV, cyan). Top rows show individual channels and bottom rows show merged 4-channel and dual-channel images to visualize co-localization of CB, CR, or PV with reporter+ neurons for (**f**) ZE1001 h56D-bReaChES-tdT, (**g**) AiP12068 DLX2.0-ChrimsonR-tdT, and (**h**) AiP14858 DLX2.0-ChR2-EYFP constructs respectively. Scale bars, 20 μm.

In this study we report optogenetically-evoked activity from seven macaques (n=30 injections across 5 brain areas, Fig. S3, S4). We characterized three viral constructs based on the “pan-inhibitory” mDLX or h56D enhancers, sometimes also called hDLXI56i (see Fig. S1d for details on the overlap and differences between enhancers)^10^. Viral constructs were created by combining a viral serotype, enhancer element variant, opsin, and reporter protein. We did not exhaustively vary each of these parameters and instead focused on viral constructs previously characterized extensively in rodents^10^. Two of the constructs that were developed at the Allen Institute contain the ‘DLX2.0’ enhancer sequence, which consists of a 3x concatenated core (overlapping region between hDlx and mDlx, see Fig. S1) region of the DLXI5/6i enhancer^8,10^. One version of this DLX2.0 construct, AAV/PHP.eB-DLX2.0-ChR2-EYFP (AiP14858)^10^, is designed to drive expression of ChR2-EYFP fusion protein, while the other variant, AAV/PHP.eB DLX2.0-ChrimsonR-tdT (AiP12068)^10^, is designed to drive expression of the red-shifted ChrimsonR opsin and tdTomato fusion protein. The third construct tested, AAV1-h56D-bReaCHES-tdTomato^9^ (ZE1001), contains a variant of the h56D enhancer and utilizes the red-shifted opsin bReaChES. Details for each construct and where they were used for *in vivo* and *in vitro* experiments are provided in the supplement (see Fig. S3).

### 2.2. Pan-inhibitory enhancers restrict reporter protein expression to GABAergic (GAD67+) neurons and preferentially target PV+ neurons

To evaluate specificity (e.g., the proportion of all neurons expressing a viral reporter that also colabel GABAergic neurons) and completeness (e.g., proportion of the total GABAergic population that shows colabeled reporter expression) of transgene expression in interneurons from these constructs, we quantified reporter protein (EYFP or tdT) positive neurons and their co-expression of immuno-labeled markers for GABAergic neurons (Fig. 1, 2). We imaged reporter protein and immunolabeled neurons across the depth of the cortex in 3 serial 60-µm thick coronal sections, sampled through the rostro-caudal extent of the injection site (Fig. 1, 2, S5). For each section we imaged two regions of interest (ROIs): one within the densest area and one in the sparsest area of the reporter protein label (Fig. 2, S5). Thus there are a total of 6 columnar ROIs (at 1272 µm wide for each column) per case.

**Figure 2:**
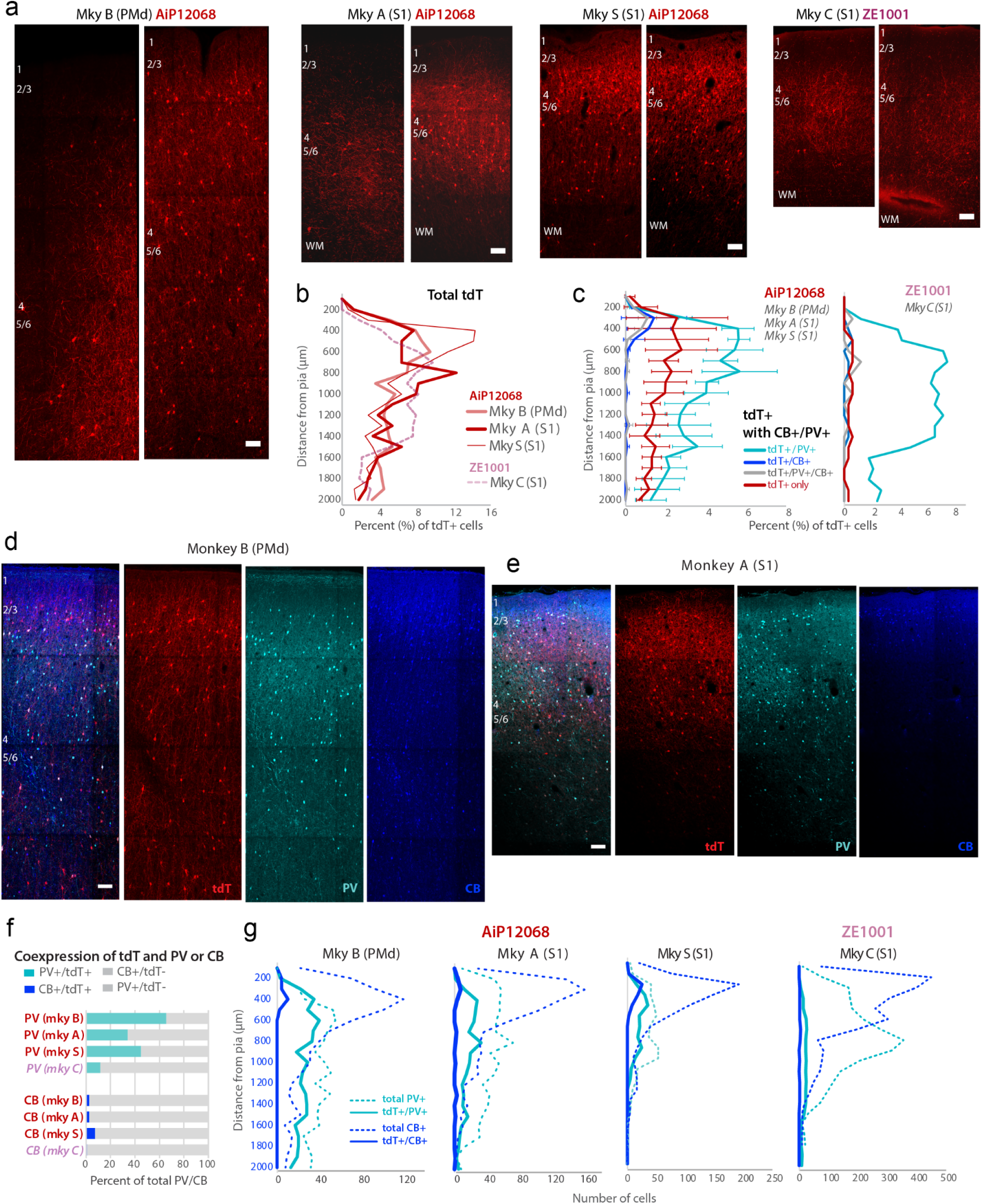
Virally-labeled cells recapitulate canonical laminar distribution of inhibitory neuron subtypes and most completely label PV+ neurons as compared to other inhibitory subpopulations across macaque cortical areas. (**a**) Representative confocal microscopy z-projection images showing tdT (red) expression along the cortical depth in coronal sections through the transduction sites of AiP12068 (Mky B, PMdc/M1; Mky A, S1; Mky S, S1) and ZE1001 (Mky C, S1). Labels mark the top of each laminar compartment (layers 1, 2/3, 4, 5/6) and the white matter (WM) boundary. (**b**) Percentage of tdT+ neurons per case (AiP12068: Mky B, PMdc/M1; Mky A, S1; Mky S, S1; shades of red and Z1001: Mky C, S1; pink dashed line) as a function of distance from the pial surface (μm). (**c**) Laminar distribution of percentage of tdT+ cells co-expressing PV (tdT+/PV+), CB (tdT+/CB+), both markers (tdT+/PV+/CB+), or neither (tdT+ only) for the AiP12068-injected monkeys (*left*; averaged n = 3 animals) and the ZE1001-injected monkey (*right*; Mky C, S1). (**d**, **e**) Representative immunofluorescence images showing merged and individual channels (tdT, PV, and CB) with cortical layer annotations for (**d**) Monkey B (PMd) and (**e**) Monkey A (S1). (**f**) Bar graphs showing the percentage of total PV+ or CB+ cells co-expressing tdT (i.e., completeness of subpopulation expression) across all animals for each viral construct. (**g**) Absolute cell counts of total PV+, tdT+/PV+, total CB+, and tdT+/CB+ cells as a function of cortical depth for all four monkeys and the AiP12068 and ZE1001 constructs. Scale bars, 100 μm.

We assessed co-expression of reporter+ somata (visualized directly without immunohistochemistry) with a pan-GABAergic marker, Glutamate Decarboxylase 67 kDa (GAD67), the enzyme essential for GABA synthesis (Fig. 1b). We found that among all the reporter+ somata, 97% of the DLX2.0 construct+ (i.e., AiP12068 and AiP14858) neurons and 81% of the h56D+ (i.e., ZE1001) neurons were also GAD67+, confirming construct specificity for GABAergic neurons (Fig. 1c). We next assessed the completeness of labeling for the total GAD67+ population and found that the AiP12068 construct labeled ∼64% of all GAD67+ neurons. However, AiP14858 and ZE1001 only labeled ∼37% and ∼32% of all GAD67+ cells respectively (Fig. 1d). Previously similar constructs were shown to express more completely in GABAergic cells in NHPs. However the viral constructs were only tested in marmoset visual cortex and only included a reporter, not an opsin-reporter fusion protein^8,9,13^.

We then assessed whether these three viral constructs were expressed in a variety of different inhibitory neuron types or preferentially expressed in specific subpopulations of interneurons over others. Reporter+ inhibitory neurons were assessed for co-expression of immunolabeled calcium-binding proteins, Parvalbumin (PV), Calbindin (CB) and Calretinin (CR) (Fig. 1e-h), which label three largely non-overlapping neurochemical populations in the macaque cortex^35^. Overall, across the three constructs we tested, we found that the majority of the reporter+ inhibitory neurons expressed PV (66-90%), with a small proportion (2-9%) expressing CB (Fig. 1e-h). No reporter+ inhibitory neuron was found to co-express CR (Fig. 1e-h). The preferential transgene expression in PV+ inhibitory neurons was found across all 3 constructs analyzed regardless of whether the injection was made in PMdc/M1 or S1 (Monkeys A, S, C, B Fig. 1e-h).

However, the degree of specificity for expressing in PV+ neurons varied by construct. For the ZE1001 (h56D) construct, ∼90% of all transgene-expressing neurons were PV+, with 4% PV+/CB+ and only 2% CB+ (Fig. 1e top, f). In contrast for both DLX2.0 constructs AiP12068 and AiP14858, ∼66-73% of the transgene-expressing neurons were PV+, and 5-9% were CB+ (Fig. 1e middle and bottom, g, h). Interestingly for these DLX2.0 constructs, ∼27% of tdT+ AiP12068 and 18% of EYFP+ AiP14858 transgene-expressing neurons lacked detectable immunoreactivity for the examined calcium-binding proteins (i.e., PV, CB, and CR) and only expressed the reporter protein (i.e., tdT+ only or EYFP+ only, Fig. 1e middle and bottom). Given that ∼97% of the reporter+ neurons from DLX2.0 constructs were GABAergic (GAD67+), we suspect that this population that only expresses the reporter could represent another unlabeled neurochemical inhibitory neuron subpopulation. Another possibility is that this is a result of virally-mediated downregulation of calcium binding proteins, as seen with rabies viral constructs^13^. Reporter+ only expression was lowest for the ZE1001 construct (4%, Fig. 1e, top), however note that this construct labels a smaller fraction (∼32%) of the total GABAergic population compared to AiP12068 (Fig. 1d).

Notably, the different constructs labeled different parts of the cell. AiP12068 (ChrimsonR-tdT) and ZE1001 (bReaChES-tdT) labeled both neuronal processes and somata (Fig. 1f, g). In contrast, AiP14858 (ChR2-EYFP) strongly labeled neuronal processes and weakly labeled few somata (Fig. 1h), likely reflecting differences in how much protein accumulates in the secretory pathway. We counted a total of n = 403-711 tdT+ somata from each case with either AiP12068 (Monkey (Mky) B n = 711, Mky S n = 619, Mky A n = 407) or ZE1001 (Mky C n = 403) injections, but only a total of 120 EYFP+ somata from the case with the AiP14858 injection (Mky B S1). Thus, the tdT+ somata were at least 3x more numerous than the EYFP+ somata in our dataset.

### 2.3. Laminar distribution of transgene expression align with layer-specific subtypes

We next analyzed the laminar specificity of transgene expression from these constructs by quantifying the number and proportion of tdT+-labeled cells across cortical depth within each of the ROIs (Fig. 2a). In all 4 cases analyzed (3 from AiP12068, 1 from ZE1001), there was a peak in tdT+ cell expression in layer 3, ∼300-800 µm from the pial surface (Fig. 2b). This pattern emerged mainly due to the population of tdT+ inhibitory neurons that co-expressed PV, and to a lesser extent from cells that were tdT+ but did not express any of the assessed calcium-binding proteins (i.e., PV, CB, and CR) (Fig. 2c). Laminar specificity for AiP14858 could not be performed as too few somata (n=120, Monkey B S1, Fig. 1h) were labeled by the construct.

We then determined the completeness of expression in inhibitory subpopulations of these viral constructs. We quantified the total number of immuno-labeled PV+ and CB+ cells, and calculated the proportion colabeled with tdT+ across the cortical depth (Fig. 2d-g). The CR+ population was not examined as no CR+ neurons were colabeled as tdT+. Our results revealed that the proportion of PV+ inhibitory neurons expressing tdT was significantly greater than that of CB+ cells (Student *t*-*test*, *p* = 0.009; Fig. 2f). The expression of tdT within the CB+ or PV+ inhibitory neuron population also varied among cases and constructs (Fig. 2f, cyan). For the AiP12068 DLX2.0 construct, ∼35-65% of the total PV+ population, and ∼3-8% of the total CB+ neurons were labeled for tdT (Fig. 2f, monkeys B, A, S). In contrast, the ZE1001 h56D construct labeled only ∼12% of the total PV+ and < 1% of the CB+ population (Fig. 2f, monkey C).

The cortical depth profiles of these transgene-expressing PV+ or CB+ inhibitory neurons (Fig. 2g), revealed specificity in laminar depth location based on neurochemical expression. Consistent with previous studies^36^, CB+ neurons were mostly concentrated within upper layers 2 to 3a, while PV+ neurons were found across all layers. PV+ neuron distribution peaked in middle layers 3b - 5a, ∼400-1000 µm from the pia in both PMd and S1 (Fig. 2g, dotted lines). This neurochemical expression-specific depth distribution was also found for the subset co-labeled as tdT+. tdT+/CB+ colabeled neurons only present in layer 2. While tdT+/PV+ colabeled neurons were located across all depths, peaking in layer 3 - 4, ∼400-500 µm from the pia (Fig. 2g, solid lines). Overall, these data demonstrate that these putatively pan-inhibitory constructs more completely express in the PV+ subpopulation of inhibitory neurons, while only modestly expressing in CB+ neurons in macaque PMd and S1 cortices.

### 2.4. Robust in vivo optotagging of inhibitory neurons in macaque cortex

Immunohistochemistry analysis of reporter expression confirmed that the putatively pan-inhibitory constructs expressed in inhibitory neurons, primarily PV+, and thus drove expression of opsins in these cells. To assess whether transgene-expressing neurons remained functionally integrated in cortical circuits, we tested whether optical stimulation could induce spiking in inhibitory neurons *in vivo* and whether connected neurons showed suppression of activity.

In a procedure under propofol anesthesia, we performed a craniotomy and reflected the dura. We then identified hotspots by visualization of reporter expression (Xite fluorescence LED, NIGHTSEA, Morgantown, PA), and slowly inserted a 32 channel Plexon V-probe into these locations. We then optically stimulated using a 473 nm laser (blue) for the ChR2 opsin and 561 nm (green)/ 638 nm (red) laser for the ChrimsonR opsin and monitored spiking activity during laser stimulation. Once we confirmed optically-evoked spiking activity at a hotspot, we adjusted the depth of the probe to maximize optically-evoked spiking activity. We then ran multiple blocks of optotagging trials at each site, and saved broadband neural activity for offline analysis.

Offline, we first used Kilosort4^34^ to identify single neurons and then further analyzed the responses of these neurons. We determined that a cell was optotagged if it responded within 10 ms of laser onset using the stimulus-associated latency test (SALT)^3^. We classified a cell as being suppressed if there was an over 50% decrease in its firing rate during the stimulation period as compared to the baseline firing rate^3^. Suppressed cells are not directly suppressed by the light stimulus, but are likely suppressed by nearby activated inhibitory neurons. Fig. 3a-c show representative examples of optotagged (Fig. 3a, b, c (bottom)) and a suppressed unit (Fig. 3c, top). Overall, *in vivo*, we optotagged 66 inhibitory units and suppressed 20 units across two brain areas (46 from S1, 40 from PMd/M1, all 3 constructs, n=7 monkeys). Typically, optotagged units demonstrated a sharp increase in firing rate, with a low latency onset (Resp. lat., ∼5 ms) and only modest adaptation (Adap. index) during the stimulation period (Fig. 3d-f). Furthermore, average spontaneous and evoked waveforms from optotagged units demonstrated a very high median correlation (>.98) across all viral constructs (Wf. corr., Fig. 3d-f).

**Figure 3:**
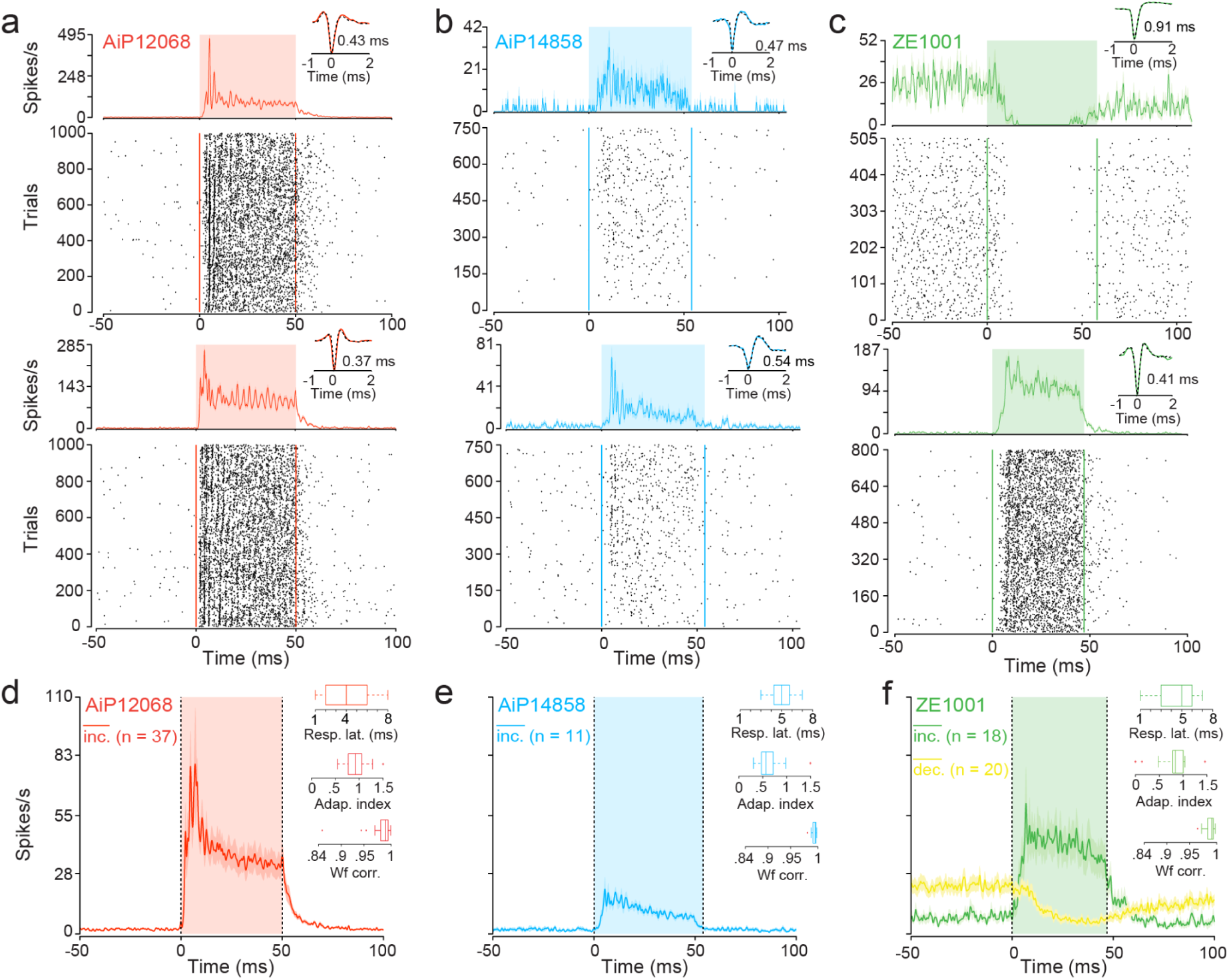
Cortical inhibitory neurons targeted via pan-inhibitory constructs robustly respond to light stimulation. (**a**, **b**, **c**) Raster plot of spike times in trials (i.e., laser pulses) and peristimulus time histograms aligned to the onset of laser pulses for representative optotagged (i.e., directly activated units that respond rapidly to laser onset (*<* 10 ms)) or a suppressed unit from three pan-inhibitory constructs. (**a**) The units for the AiP12068 and (**c**, *bottom*) a unit for the ZE1001 constructs increase in firing rate robustly whereas (**b**) the units for the AiP14858 construct only moderately increase in firing rate as a function of laser onset. (**c**, *top*) Exemplary suppressed unit that decreased in firing rate at laser onset for the ZE1001 construct, likely due to activation of inhibitory neurons. *Insets*: The average evoked (solid color) and spontaneous (dotted black line) waveforms from spike times in the peri-stimulus time period. (**d**, **e**, **f**) Average peristimulus time histograms aligned to laser onset for each of the three pan-inhibitory constructs (AiP12068, AiP14858, ZE1001) from units pooled across all seven monkeys and two cortical brain regions (i.e., PMdc/M1 & S1). Units for AiP12068, AiP14858, and ZE1001 increased (inc., red, blue, and green respectively) and only for ZE1001 decreased (dec., yellow-green) in firing rate. n is the number of units per construct that either increased or decreased in firing rate. *Insets*: For all increase units per construct: boxplots of response latency (Resp. lat.), when a unit first reaches ½ peak firing rate post-laser onset; adaptation index (Adap. index), ratio of mean firing rate for a unit in late as compared to early stimulation period; waveform correlation (Wf. corr.), correlation between the average spontaneous waveform for a unit and its average evoked waveform.

To assess which viral construct led to the strongest modulation, we averaged the firing rates of optotagged units aligned to laser onset grouped by viral construct. A Kruskal-Wallis test revealed that there was a main effect of construct on mean firing rate during the stimulation period (H(2) = 10.43 *p* = 0.0054). Units expressing constructs carrying red-shifted opsins, AiP12068 and ZE1001, robustly and significantly elicited the largest median firing rate during stimulation across 37 units (Fig. 3d) and 18 units (Fig. 3f), respectively. In contrast, in units expressing a blue-sensitive opsin, AiP14858, stimulation only elicited a moderate increase in firing rate (11 units; Fig. 3e). Post-hoc tests further confirmed the pattern (Wilcoxon rank-sum with Bonferroni corrected p-values, AiP12068 (median ± interquartile range: 20.87 ± 36.89 spikes/s) and AiP14858 (10.64 ± 4.81 spikes/s) *p* = 0.011; AiP14858 and ZE1001 (30.44 ± 30.23 spikes/s) *p* = 0.013; AiP12068 and ZE1001 *p* = 1). Finally, for units expressing the transgene of the ZE1001 construct, 18 cells increased their firing rate and 20 cells decreased their firing rate during the stimulation period (Fig. 3f).

### 2.5. Multidimensional clustering algorithm reveals inhibitory neurons targeted by pan-inhibitory constructs

Our immunohistochemistry data indicate that most neurons labeled by the DLX5/6 enhancer-based constructs were, of the calcium-binding proteins examined, PV+, with a minority of the neurons expressing CB (typically thought to be a subpopulation of SST neurons). Mouse and primate studies suggest that PV+ neurons often have the narrowest action potential waveforms, whereas SST+ neurons have intermediate widths^1,3^. Based on these results and our immunohistochemistry data, we predicted that optotagged neurons would be predominantly narrow-spiking with a smaller proportion of intermediate-width waveforms. Second, PV+ neurons preferentially synapse onto excitatory neurons^37^, which typically have large waveform widths^1,38^. Accordingly, we predicted that neurons that suppress their activity in response to laser stimulation would largely have broader waveforms.

We tested these two predictions by analyzing the waveforms from our *in vivo* experiments. We averaged the peri-stimulus waveforms for each of the 86 units (directly optotagged and suppressed), normalized them to between a range of -1 to +1, and polarity-corrected them to be negative spiking units (so as to avoid clustering based on polarity). These average waveform shapes were entered into WaveMAP^25^, our algorithm for classifying cell types based on the overall waveform shape.

The results of our WaveMAP analysis are shown in Fig. 4a. Each unit is represented as a circle whose diameter reflects trough-to-peak duration (ms). Green and yellow colors indicate whether a unit increased or decreased its firing rate, respectively (Fig. 4a). This analysis qualitatively revealed two distinct clusters which separate the ‘optotagged’ (green) units from the ‘suppressed’ (yellow) units. The waveforms segregate along the UMAP 1-axis with the narrowest, optotagged waveforms on the left to the widest, suppressed waveforms on the right (Fig. 4a). The cluster of suppressed waveforms on the right is likely to be enriched for excitatory cells as they tend to have the widest waveforms and decrease in firing rate in response to laser onset (Fig. 4a).

**Figure 4:**
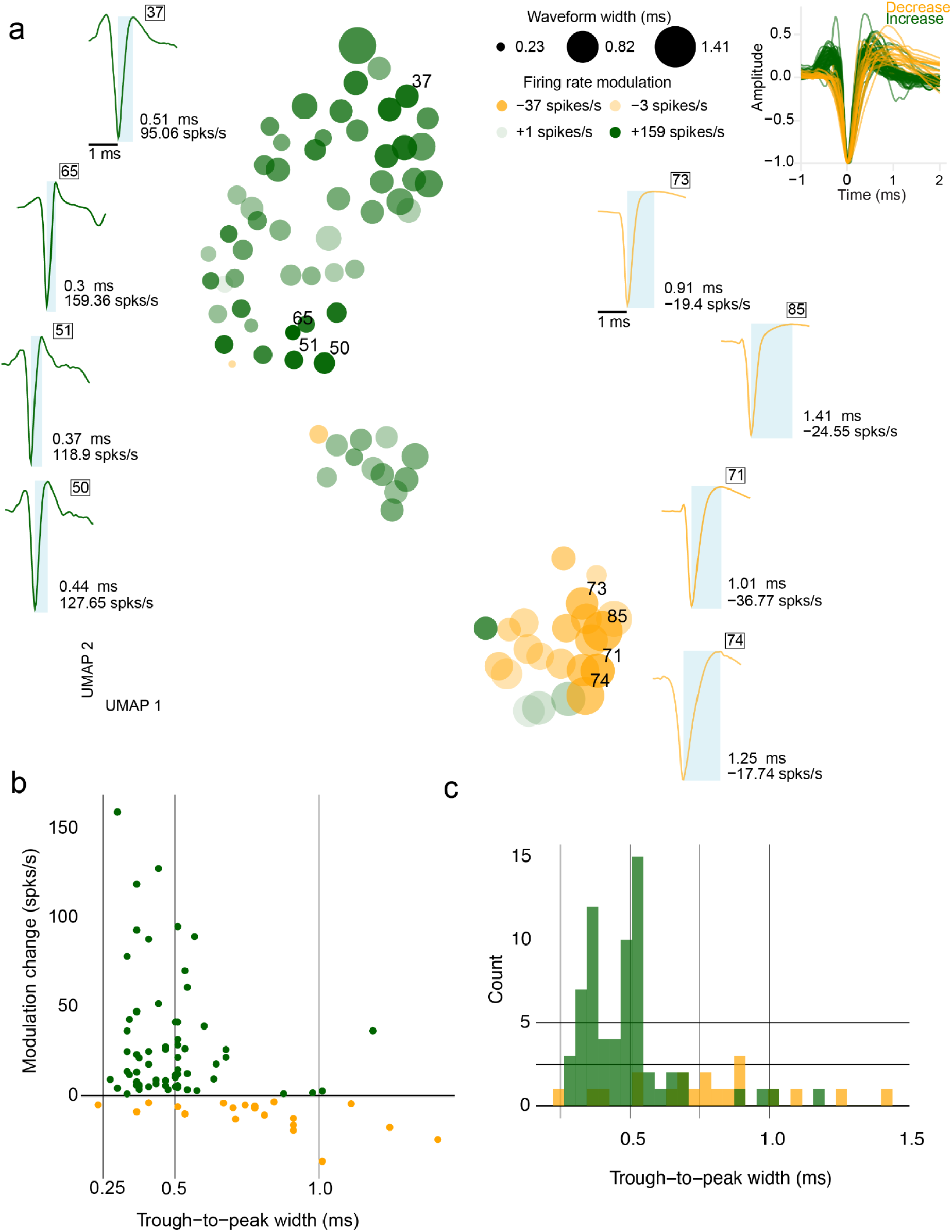
Optotagged cortical inhibitory neurons exhibit narrow waveforms and cluster separately from putatively excitatory neurons. (**a**) UMAP plot of n = 86 average unit waveforms (amplitude-normalized and polarity matched to be negative) pooled from 7 monkeys, 2 cortical areas (PMdc/M1 & S1) and 3 pan-inhibitory constructs (AiP12068, AiP14858, and ZE1001). Units are represented as circles of varying width, indicative of waveform width (i.e., 0.23, 0.82, and 1.41 ms), and color, indicating the strength (opacity) of either an increase (green) or decrease (yellow) in firing rate in response to laser pulses. Individual waveforms with trough-to-peak duration, average difference from baseline firing rate (spks/s), and unit number are represented around the UMAP plot. *Inset*: All trough-aligned and amplitude-normalized waveforms included in the WaveMAP visualization. (**b**) Scatterplot of waveform width (trough-to-peak width) by the change in firing rate from baseline (modulation change). Green and yellow circles increased and decreased in firing rate respectively. (**c**) Histogram of trough-to-peak width demonstrating the distribution of inhibitory units generally having narrower waveforms as compared to putatively excitatory neurons (i.e., neurons with broad waveforms that decreased in firing rate).

Overall, the waveforms for cells that increased their firing after laser stimulation onset had narrower widths (Median ± interquartile range: 468 µs ± 142 µs) than those that were suppressed by laser stimulation (777 µs ± 276 µs; Wilcoxon rank-sum test comparing trough-to-peak duration: *p* = 1.85 x 10^-5^). In addition, consistent with our immunohistochemistry results showing the pan-inhibitory enhancers label both PV+ and a modicum of CB+ neurons, we observed a range of trough-to-peak widths among optotagged cells from narrow (∼.30 ms) to moderate (∼.5 ms) width (Fig. 4b, c)^3^.

### 2.6. Optotagged inhibitory neurons exhibit interneuron morphology and recapitulate characteristic biophysical membrane properties

Our *in vivo* optotagging experiments confirmed that transduced neurons can express functional opsins and can elicit robust spiking. We next used *in vitro* whole-cell patch clamp recordings to further validate opsin expression in transduced neurons. *In-vitro* physiology offers a complementary and more direct functional assessment of labeled optotagged neurons, allowing us to characterize intrinsic membrane properties and morphological features, and to compare kinetics of optically-versus current-evoked action potentials.

We first focused on tdT+ cells in acute slices harvested after injections of AiP12068 (Fig. 5a). These experiments revealed that recorded tdT+/ChrimsonR+ cells were light responsive (n = 4) with action potential firing responses time-locked to 10 ms, 565 nm light pulses at 0.6-3 mW power. Increasing light pulse duration from 50, 100, to 200 ms elicited trains of action potentials with an increasing number of spikes (Fig. 5b, c). These data confirmed the functional optotagging of these neurons, based on their expression of ChrimsonR opsin.

**Figure 5:**
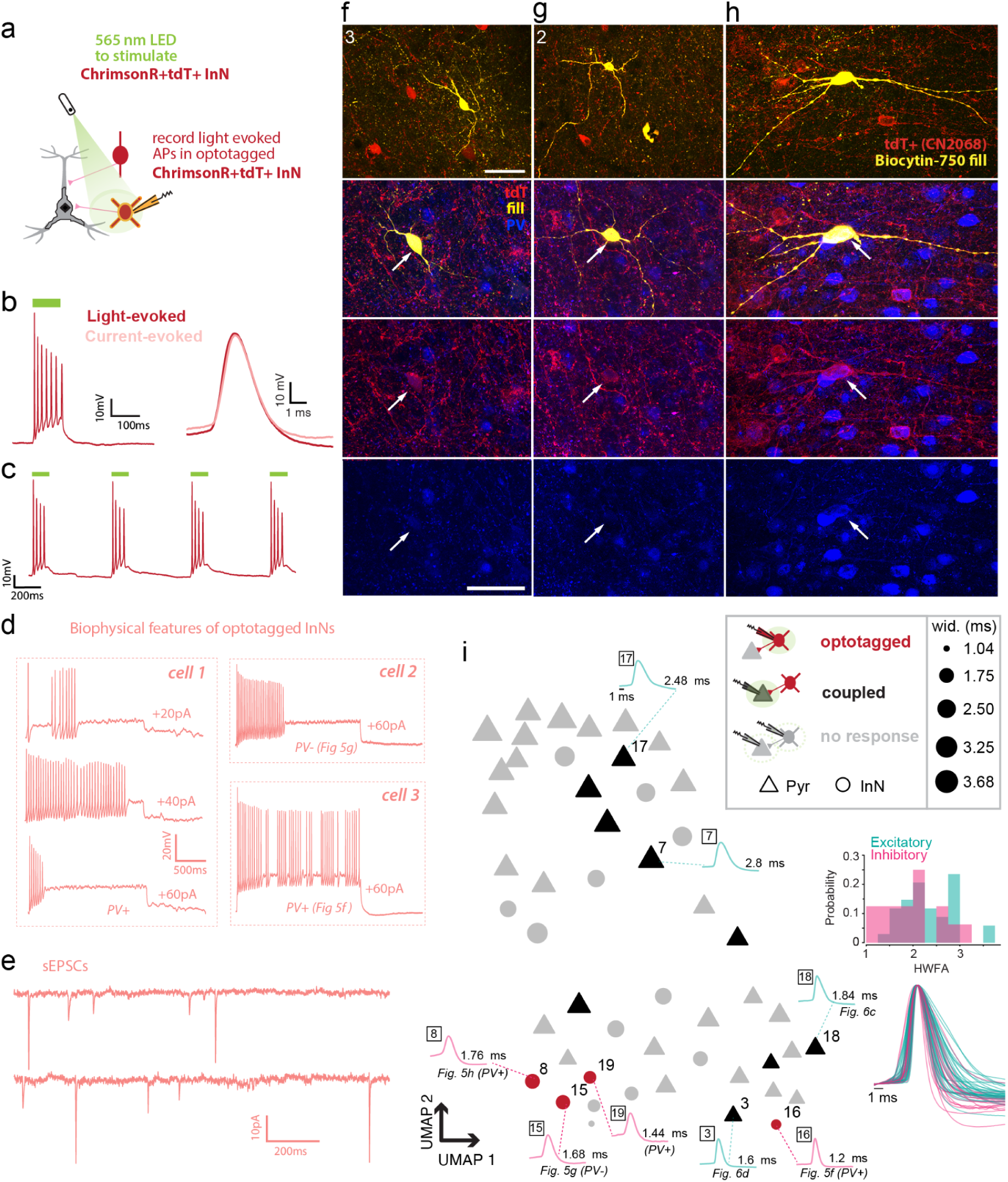
Optotagged inhibitory neurons exhibit classic interneuron morphology and similar optically-evoked and current-evoked waveforms. (**a**) Schematic showing *in vitro* whole-cell patch clamp recording and LED light stimulation of visually identified ChrimsonR+/tdT+ inhibitory interneuron (InN) expressing the CN2068 construct. Interneurons were identified based on their round somata appearance in IR-DIC optics as well as presence of tdT+ label. (**b**) Whole-cell patch clamp recording from a tdT+ interneuron in Monkey A. *Left* shows an optically-evoked train of action potentials (APs) recorded during a 100 ms 565 nm LED light pulse. *Right* shows a single optically-evoked action potential from this train (3rd action potential), expanded to visualize waveform kinetics superimposed onto the equivalent action potential waveform evoked by current injection, in the same cell. Note that the light vs current-evoked action potential waveforms have the same kinetics. (**c**) Action potential train responses to a series of 100 ms light pulses, confirm that action potential trains time lock to optical stimuli. (**d**) Representative voltage traces of repetitive current-evoked action potential firing responses to 2 s current injections of 3 optotagged neurons. Responses of one neuron (cell 1) to increasing current injections shows heterogeneity in firing pattern: phasic response to +20 pA, sustained high frequency response to +40 pA, fast-adapting response to +60 pA. Two neurons (cells 2 and 3) show distinct firing patterns in response to +60 pA current injections: Note the fast-spiking, fast-adapting phenotype of a PV-/Tdt+ neuron (cell 2, same cell shown in **g**) and the fast-spiking stuttering firing pattern of a PV+/tdT+ neuron (cell 3, shown in **f**). PV+/-status confirmed via post-hoc immunolabeling. (**e**) Representative traces of spontaneous excitatory postsynaptic currents (sEPSCs, Vhold -80mV) recorded from an optotagged PV+ inhibitory neuron showing characteristic fast kinetics of individual synaptic events. (**f, g h**) Confocal Z-max projection images of tdT+ cells expressing the CN2068 transgene (red) that were recorded and filled with biocytin and visualized with Streptavidin 750 (yellow), and subsequently immunolabeled for PV (blue). Row 1: Dual channel low magnification images of the fills reveal multipolar morphology of CN2068-expressing tdT+ interneurons. Rows 2-4: Merged triple channel and single channel z-projection image stack (5-optical planes) of somata showing colocalization of fills (yellow) with tdT+ (red) and PV+ (blue) labeling. PV+ cells are shown in **f** and **h**; and a PV-cell is shown in **g**. Scale bar = 50µm. (**i**) UMAP plot of n = 50 cell waveforms (amplitude-normalized) pooled from 9 monkeys and 8 cortical areas. Cells were included for this analysis as they were identified as coupled or optotagged (CN2068, CN4858; n=13) or when a biocytin-filled cell from a previous recording was visually identified as having either a pyramidal (excitatory) or interneuron (inhibitory) morphology (n=37). Cells are represented as triangles (excitatory pyramidal neurons) or circles (inhibitory interneurons) of varying width (half width full amplitude duration; 1.04 -3.68 ms) and color (black -coupled, red -optotagged, gray -no response). Individual waveforms with half-width full amplitude (HWFA) duration and cell number are plotted around the UMAP plot. In addition all waveforms included in the UMAP visualization and a histogram of the HWFAs of the waveforms are plotted to the right (Excitatory -cyan, Inhibitory -pink).

To further characterize the functionality of these visually and functionally validated AiP12068 optotagged cells, we measured their intrinsic passive and active membrane properties in response to a series of hyperpolarizing and depolarizing current steps (Fig. 5d, e). We found that these cells have biophysical properties consistent with GABAergic interneurons, with high excitability (reviewed in^39^). Specifically, these AiP12068 optotagged cells on average have high input resistance, low rheobase, and narrower current-evoked action potential waveforms (Fig. S6b, c). The cells also had a prominent sag potential and fast after-hyperpolarization (Fig. S6b, c), indicating the enrichment of HCN and K channels^40–44^.

The kinetics of these current-evoked action potential responses in optotagged tdT+ cells were consistent with the waveform and repetitive firing patterns of optically-evoked action potential responses (see Fig. 5b). Trains of action potentials show an increase in firing frequency in response to increased current step amplitude (see Fig. 5d and Fig. S6a). All cells exhibited heterogeneous high frequency firing patterns. One cell showed different firing patterns in response to increasing current injections, with phasic, sustained high frequency and fast-adapting responses to +20pA, +40pA and +60pA, respectively (Fig. 5d, cell 1). Two cells exhibited fast-adapting, high frequency action potentials in response to higher current steps (Fig. 5d cell 2 and Fig. S6a (top MCC)). One cell exhibited a high frequency stuttering firing pattern^45^ (Fig. 5d, cell 3). In addition to membrane properties, recordings under voltage clamp (Vhold -80mV) revealed that the majority of the spontaneous excitatory postsynaptic currents were fast, a phenotype typically associated with GABAergic inhibitory interneurons^41^ (Fig. 5e, Fig. S6d).

Intracellular filling and confocal imaging of recorded tdT+ neurons confirmed that they were non-pyramidal and aspiny in their morphology (Fig. 5f-h). Filled neurons were also immunolabeled to determine PV+ expression. The neuron that showed phasic, sustained, and fast-adapting action potentials, depending on the current step (Fig. 5d cell 1), was confirmed to be multipolar in morphology (i.e., having numerous primary dendritic branches that extended vertically, tangential to the pial surface) and PV+ (confocal imaging not shown). The stuttering neuron was confirmed to be vertically oriented, multipolar in morphology, and was moderately PV+ (Fig. 5d cell 3, f). For the two fast-adapting optotagged neurons, one was a small multipolar neuron that was PV-(Fig. 5d cell 2, g) and the other was a large multipolar neuron and was PV+ (Fig. 5h, Fig. S6a (top MCC)).

In order to assess how actional potential waveform properties of optotagged inhibitory neurons compared to other neurons, *in vitro* waveform data were entered into a WaveMAP analysis (Fig. 5i). 50 neurons were morphologically-identified as either layer 3 pyramidal excitatory neurons or as inhibitory neurons (Table 2 details the source of the neurons from the current cases and archived database; 8 cortical areas from 9 monkeys). This dataset also included waveforms from neurons that were synaptically coupled with optogenetically-labeled fibers from 2 cases injected with either AiP12068 and AiP14858 in the current study (Fig. 5i). This analysis qualitatively reveals two distinct clusters which separate neurons with narrower waveforms (bottom right) from those with wider waveforms (upper left, Fig. 5i). Most of the inhibitory neurons have narrower waveforms and appear in the bottom right cluster, consistent with PV+ neurons (all optotagged neurons appear in this cluster). While the remaining inhibitory neurons in the upper left cluster demonstrate wider waveforms, consistent with SST+ neurons^3^. We found that on average inhibitory neurons had narrower waveforms (median ± interquartile range: 1.92 ± 0.88 ms) than excitatory neurons (2.32 ± 0.96 ms). However this difference was not significant (Wilcoxon rank-sum; *p* = .094). This is likely due to the analysis being underpowered as there are only 16 inhibitory neurons included in this analysis (Fig. 5i, histogram and stacked waveforms).

**Table 1.** Primary and secondary antibodies used to perform immunohistochemistry.

| **Primary Antibody** | **Vendor** | **Catalog #** | **RRID #** | **Dilution** |
|---|---|---|---|---|
| rabbit anti-CR | Swant | CR7697 | AB_2619710 | 1:500 |
| goat anti-CB | ThermoFisher | PA5-46936 | AB_2609696 | 1:100 |
| guinea pig anti-VGAT | Synaptic Systems | 131004 | AB_887873 | 1:400 |
| mouse anti-PV | Swant | 235 | AB_10000343 | 1:1000 |
| guinea pig anti-IBA1 | Synaptic Systems | 234-308 | AB_2924932 | 1:500 |
| goat anti-GAD67 | R&D Systems | AF2086 | AB_2107724 | 1:100 |
| mouse anti-GFAP | Synaptic Systems | 173-211 | AB_2721090 | 1:500 |
| rabbit anti-cleaved caspase direct conjugate | Cell Signaling | 9664BF | AB_2070042 | 1:100 |
| **Secondary Antibody** | **Vendor** | **Catalog #** | **RRID #** | **Dilution** |
| donkey anti-goat F(ab')2 405 | Jackson | 705-476-147 | AB_2632563 | 1:200 |
| donkey anti-rabbit F(ab')2 488 | Jackson | 711-546-152 | AB_2340619 | 1:200 |
| donkey anti-guinea pig F(ab')2 647 | Jackson | 706-605-148 | AB_2340476 | 1:200 |
| donkey anti-mouse F(ab')2 750 | Abcam | 175735 | AB_3678778 | 1:200 |
| donkey anti-guinea pig F(ab')2 DyLight 405 | Jackson | 706-476-148 | AB_2632564 | 1:200 |
| donkey anti-goat IgG 488 | Jackson | 705-545-147 | AB_2336933 | 1:200 |
| donkey anti-mouse IgG 750 | Abcam | 175739 | AB_2936438 | 1:200 |

**Table 2.** All cells included in the *in vitro* WaveMAP visualization (Figure 5i) Please note that the apparent discrepancy in the number of synaptically coupled cells here (as compared to SFig 3d) is due to some coupled cells not having enough current-evoked action potentials (e.g., having one or no actional potentials) to be included in this analysis. Acronyms: ID -animal identification, nCells -number of cells, nExc -number of excitatory cells, nInh -number of inhibitory cells, nCoup -number of coupled cells, nTag -number of optotagged cells, and nPV -number of PV+ cells.

| **ID** | **Area** | **Construct** | **nCells** | **nExc** | **nInh** | **nCoup** | **nTag** | **nPV** |
|---|---|---|---|---|---|---|---|---|
| 1W | PMdr | N/A | 2 | 2 | 0 | N/A | N/A | N/A |
| 3M | S1 | 4858 | 2 | 2 | 0 | 2 | 0 | 0 |
| 6A | d46 | N/A | 2 | 2 | 0 | N/A | N/A | N/A |
| 6A | MCC | 2068 | 4 | 2 | 2 | 2 | 2 | 2 |
| 6A | dLPFC | N/A | 4 | 2 | 2 | N/A | N/A | N/A |
| 6A | vLPFC | 2068 | 7 | 5 | 2 | 5 | 2 | 1 |
| 7C | PMd | N/A | 1 | 1 | 0 | N/A | N/A | N/A |
| 7C | dACC | N/A | 1 | 1 | 0 | N/A | N/A | N/A |
| 7C | dLPFC | N/A | 2 | 2 | 0 | N/A | N/A | N/A |
| 7C | PMdr | N/A | 1 | 0 | 1 | N/A | N/A | N/A |
| 8T | PMdc | N/A | 6 | 5 | 1 | N/A | N/A | N/A |
| 9P | ACC/MCC | N/A | 8 | 6 | 2 | N/A | N/A | N/A |
| aP | A46 | N/A | 1 | 1 | 0 | N/A | N/A | N/A |
| dW | dACC | N/A | 2 | 0 | 2 | N/A | N/A | N/A |
| eN | A24 | N/A | 1 | 1 | 0 | N/A | N/A | N/A |
| eN | A9 | N/A | 1 | 1 | 0 | N/A | N/A | N/A |
| eN | dACC | N/A | 2 | 1 | 1 | N/A | N/A | N/A |
| eN | HPC | N/A | 1 | 0 | 1 | N/A | N/A | N/A |
| eN | vACC | N/A | 1 | 0 | 1 | N/A | N/A | N/A |
| eN | vA46 | N/A | 1 | 0 | 1 | N/A | N/A | N/A |
|  |  | **Totals** | 50 | 34 | 16 | 9 | 4 | 3 |

### 2.7. Transgene-expressing interneurons elicit optically-evoked inhibitory postsynaptic currents in coupled postsynaptic neurons

We further validated the functionality of inhibitory synapses of transgene-expressing interneurons by optically-evoking inhibitory synaptic currents in coupled neurons. Thereby demonstrating that transgene-expressing interneurons remain functionally integrated in local circuits (Fig. 6).

**Figure 6:**
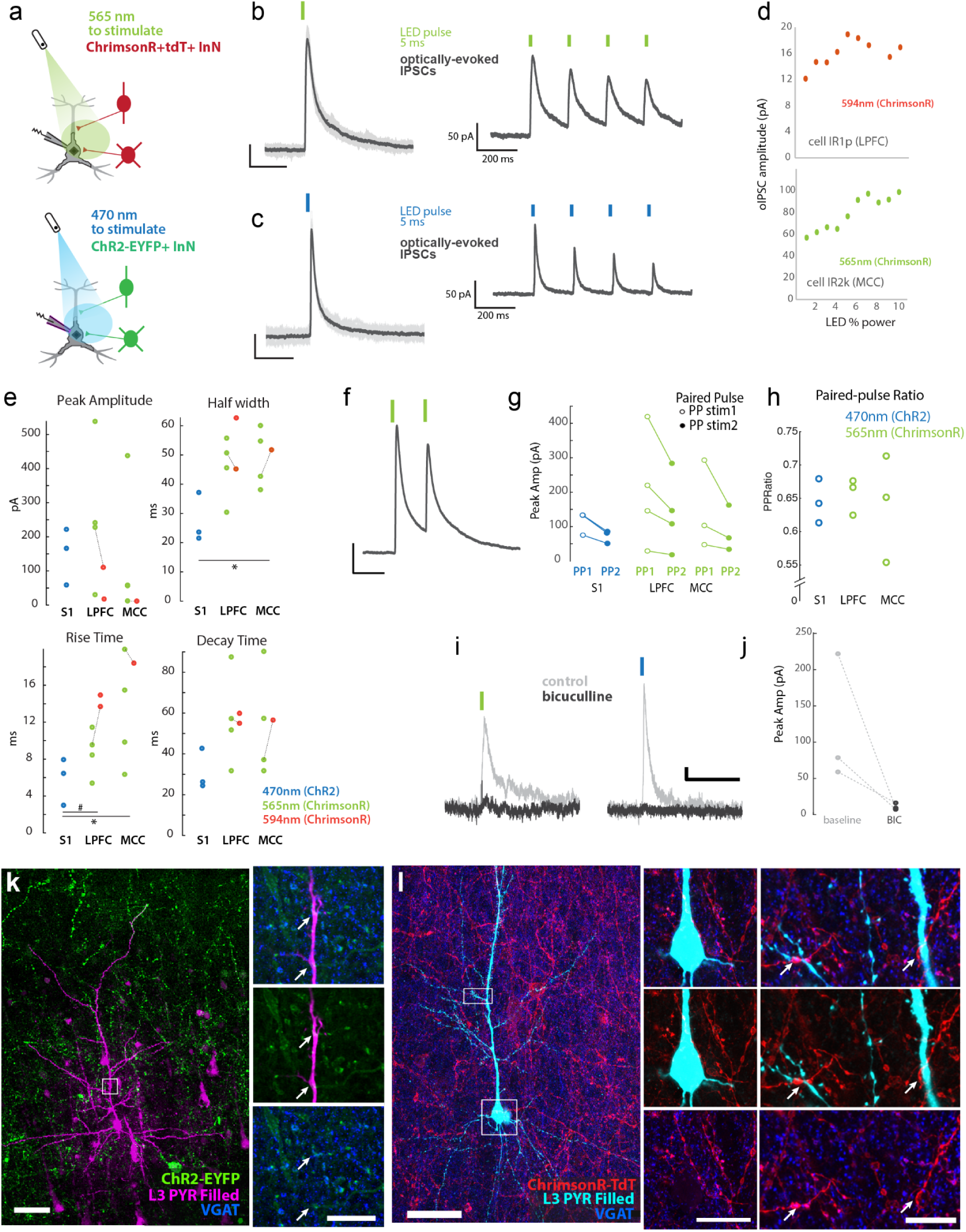
Functional, transgene-expressing, GABAergic fibers optically evoked inhibitory currents in synaptically coupled pyramidal neurons. (**a**) Schematic showing *in vitro* whole-cell patch clamp recording in pyramidal neurons synaptically-coupled to inhibitory neurons expressing ChrimsonR/tdT via AiP12068 or ChR2/EYFP via AiP14858. (**b, c**) Optically-evoked inhibitory postsynaptic currents (oIPSCs) in response to a single and a train of 5 ms LED pulses: 565 nm for ChrimsonR (**b**, green bar); 470 nm for ChR2 (**c**, blue bar). (**d**) Examples of input-output relationships (LED power x oIPSC amplitude) for representative cells recorded and stimulated with 2 wavelengths (collimated LED; 594 and 565 nm). Responses did not depend on wavelength and were saturated at low power for most cells. (**e**) Scatter plot of oIPSC peak amplitude and kinetic properties (i.e., half width, rise time, decay time) measured at the first maximal response stimulus power, by opsin, light stimulus wavelength (470 nm for ChR2, 565 or 594 nm for ChrimsonR) and area. Dotted lines connect data points from the same neuron stimulated by two wavelengths. (**f**) Representative current traces showing paired-pulse oIPSC responses. (**g**) Peak oIPSC amplitudes to paired-pulse stimulation (PP stim; 200 ms apart) and (**h**) paired-pulse ratios [peak 2/peak 1] for individually recorded cells. (**i**) Representative control and post bicuculline application oIPSC responses for the ChrimsonR (green bar) and ChR2 (blue bar) constructs. (**j**) Scatter plots showing baseline oIPSCs and oIPSCs 5 mins post bath application of GABA blocker, bicuculline (10 µM). Bicuculline (BIC) diminished oIPSCs by 70-95%. (**k, l**) Confocal images of pyramidal neurons (L3 PYR) filled with biocytin during recording (magenta in **k**, cyan in **l**), coupled to ChR2+ (k, EYFP green) or ChrimsonR+ inhibitory interneurons (l, tdT red). *Insets*: (**k**, l *right* panels) are higher magnification to show appositions of filled dendritic segments with reporter+ fibers co-labeled with VGAT (blue), a marker for inhibitory synapses. Scale bar = 20µm.

In order to assess the synaptic coupling of layer 3 (L3) pyramidal neurons with light-sensitive inhibitory axon terminals, we recorded from unlabeled L3 pyramidal neurons located within a plexus of either ChrimsonR-tdT+ (i.e., AiP12068+ neurons) or ChR2-EYFP+ fibers (i.e., AiP14858+ neurons) (Fig. 6a, b, c, e, k, l). Synaptic coupling was characterized by an optically-evoked inhibitory postsynaptic current (oIPSC) response (Vhold -40 mV) that was time-locked (within 100 ms) to a 5 ms light stimulus pulse (LED of 565 or 594 nm or laser of 561 nm for ChrimsonR and LED of 470 nm for ChR2). As a negative control, we confirmed the absence of the oIPSC response to the non-specific/incongruent light stimulus wavelength outside each opsin’s activation spectrum. In total we found n = 5/13 lateral prefrontal cortex (LPFC, area v46) and n = 4/10 mid-cingulate cortex (MCC, area 24c) pyramidal neurons were synaptically coupled to ChrimsonR+ interneurons (from Monkey A). Additionally we found n = 3/9 primary somatosensory cortex (S1) pyramidal neurons from Monkey M that were coupled to ChR2+ interneurons. Overall 12/32 pyramidal neurons recorded within fields of tdT+ or EYFP+ fibers in two cases (Monkey A and M) were synaptically coupled to optogenetically-labeled interneurons (Fig. 6e).

We evaluated the biophysical signaling capacity and health of these coupled pyramidal neurons by measuring their intrinsic passive membrane (Vr, tau and Rn; Fig. S6b), active membrane, and action potential firing properties (sag potential, rheobase, action potential waveform and repetitive firing properties; Fig. S6c). We confirmed the regular spiking nature^46^ of these pyramidal neurons and their ability to fire sustained repetitive trains of action potentials in response to 2 s current injections (Fig. S6a). We further confirmed the presence of robust spontaneous excitatory and inhibitory postsynaptic currents in these pyramidal neurons, indicating they received functional synaptic inputs from excitatory and inhibitory presynaptic partners (Fig. S6d-g).

Once the presence of synaptically coupled oIPSCs was confirmed, we established the input-output curve of oIPSC responses in each recorded neuron by incrementally increasing light power and measuring current peak amplitude at each stimulus intensity (1-10% LED power ∼0.65-3 mW, min power at 1% = 0.65-0.68 mW 565 nm; 0.65-1.1 mW 470 nm, 0.85-1.1 mW 591 nm) (Fig. 6d, two example neurons). 5/12 of the synaptically coupled pyramidal neurons had saturated oIPSC responses, with maximum amplitudes already evoked at the lowest possible light power setting (∼0.65 mW). The maximum oIPSC peak amplitudes recorded across pyramidal neurons were heterogenous, and did not vary by area, opsin (ChR2 vs ChrimsonR), or light stimulus wavelength (470 nm for ChR2, and 565 nm or 591 nm for ChrimsonR) and source (collimated LED vs laser fiber optic) (Fig. 6e). However, pair-wise comparisons showed a significant between-area difference in the kinetics of the oIPSCs – specifically rise time and half width duration (Fig. 6e). We also demonstrate that this optogenetic stimulation approach enables investigation of regional differences in inhibitory circuit properties. Specifically, we demonstrate heterogeneity of oIPSC responses across areas, with LPFC and S1 pyramidal neurons generally having faster oIPSCs (shorter rise times and half widths) compared to those of MCC pyramidal neurons (paired t-test p < 0.05; Fig. 6e).

To further determine the utility of these viral tools for functional circuit studies, we assessed presynaptic short term plasticity facilitation and depression of oIPSC responses (Fig. 6f-h). We evaluated paired-pulse ratios, calculated as the ratio of the peak amplitudes of oIPSCs evoked by two consecutive light pulses 200 ms apart (peak 2/peak 1). We found that for all recorded pyramidal neurons, oIPSC paired-pulse ratios were marginally < 1 (first oIPSC larger than the 2nd oIPSC), indicative of a moderately depressing synaptic connection^47^ (Fig. 6g, h). As with most oIPSC properties, no significant differences in paired-pulse ratios were found by area, opsin, or light stimulus wavelength and source (Fig. 6h).

Next, we functionally validated the GABAergic nature of these oIPSCs from transgene-expressing neurons using the GABAA receptor blocker bicuculline (Fig. 6i, j). For a subset of pyramidal neurons, oIPSCs were recorded before (baseline) and after 5 mins of external bath application of 10 µM bicuculline. We found that bicuculline elicited a 70-95% decrease in oIPSC peak amplitude compared to baseline (Fig. 6j). Post-recording visualization of intracellularly-filled recorded pyramidal neurons and immunolabeling vesicular GABAergic transporter (VGAT) within recorded fields, confirmed that the tdT+ and EYFP+ “optotagged” boutons were VGAT+, and are therefore GABAergic (Fig. 6k, l).

Finally, past work has suggested that fluorescent reporter proteins can aggregate within target cells^48,49^ and lead to cell death. In a second series of immunohistochemistry experiments, we evaluated whether transgene expression in GABAergic neurons was associated with cell damage by assessing cleaved caspase-3 expression, a widely used marker of apoptosis and cell death^50^. We found that among all reporter+ neurons, ∼30-33% of AiP12068 tdT+ (Mky B PMdc/M1, Mky S S1), ∼57% of AiP14858 EYFP+ (Mky B S1), and ∼54% of ZE1001 tdT+ (Mky C S1) labeled neurons co-expressed cleaved caspase-3 (Fig. S5a, b). We also assessed markers of microglia (Ionized calcium-binding adaptor molecule 1 (Iba1), Fig. S5c-f) and astrogliosis (Glial Fibrillary Acidic Protein (GFAP)), within each ROI with varying densities of reporter+ cells. We found that gliosis was associated with proximity to the needle penetration but not with the density of reporter+ cells (Fig. S5c-f).

## 3. Discussion

Our long-term goal is to understand how inhibitory neurons sculpt circuit dynamics underlying cognition and behavior in the primate brain^7,15^. Towards this goal we tested three putative pan-inhibitory viral constructs in multiple areas of the rhesus macaque cortex. All three constructs were based on enhancers of the *Dlx5/6* gene, widely used regulatory elements for targeting opsin expression to GABAergic neurons in the adult mammalian cortex^8,51^. All three constructs restricted opsin expression largely to inhibitory neurons, with 81 to 97% of reporter+ labeled cells being GAD67+, and could be used to functionally optotag and modulate these cells *in vivo* and *in vitro*. Among the three, AiP12068 (DLX2.0-ChrimsonR-tdT) was most effective for *in vivo* optotagging, while ZE1001 (h56D-bReaChES-tdT) was most effective for *in vivo* suppression of neural activity. Importantly, optotagged cells remained identifiable as spontaneous waveforms were highly correlated with evoked waveforms for all 3 viral constructs. *In-vitro* patch-clamp recordings confirmed that transgene-expressing neurons retained functional GABAergic synapses capable of inhibiting synaptically coupled pyramidal neurons. Immunohistochemistry confirmed both their molecular identity and cellular health, with ChR2-EYFP and bReaChES-tdT-expressing constructs showing greater signs of toxicity than the ChrimsonR-tdT construct.

However, contrary to our expectation of transgene expression in most inhibitory neurons, the ZE1001, AiP14858, and AiP12068 transgenes expressed only in 32%, 37%, and 64%, respectively, of all GABAergic cells. Estimates for the ChR2-EYFP construct are less certain, since its reduced somatic filling made counting harder. We further found strong selectivity for PV+ neurons, with only modest labeling of CB+ neurons and no detectable labeling of CR+ neurons, consistent with prior reports in macaque V1 and rat using Dlx5/6-based constructs across multiple AAV serotypes^7,11^. Those studies established the PV preference and the scarcity of CR+ labeling, and we extend them by quantifying the intermediate position of CB+ neurons. Further, we expand on prior work, which was limited to macaque V1^7,11^, and found similar patterns of targeting of GABAergic subpopulations by these constructs across heterogeneous sensory and higher-order association areas (S1, LPFC, MCC and PMd). Importantly, we demonstrate the efficacy of these viral constructs for multimodal *in vivo* and *in vitro* functional assessments of GABAergic neurons and circuits in these diverse cortical areas in macaques.

Although previous studies have shown that these three calcium binding proteins label ∼70-80% of the total GABAergic neurons in multiple cortical areas in rhesus monkeys^52,53^, the caveat remains that our immunohistochemistry does not fully capture the diversity of inhibitory neurons^54,55^. The ZE1001 construct demonstrated the highest specificity for PV (with only 4% uniquely labeled as reporter+), although this construct only labeled ∼32% of the total GAD67+ GABAergic population. Cells carrying the transgene but negative for PV, CB and CR made up 18 and 27% of labeled neurons for the two DLX2.0 constructs. Thus we currently cannot establish what these unclassified neurons are, nor which subtypes escaped labeling altogether. Multiplexed FISH, or spatial transcriptomics with probes covering interneuron subtypes alongside the transgene, would identify both the labeled cells our markers miss and the populations these constructs fail to reach^54,55,56^. Such techniques are only now becoming turnkey for NHP tissue^54,55^ and future work will attempt to deploy such approaches to answer these questions.

Another caveat is that these specificity and completeness values are estimates, based on visual scoring of cells that expressed the reporter and the markers of interest. Weakly expressing cells are counted as negative, so completeness is likely a lower bound. How strongly a cell expresses, and therefore whether it is counted or not, can depend on intrinsic fluorescent protein-brightness^48^, fusion versus separated expression, and soma targeting. Our completeness and specificity estimates used the opsin-bearing constructs themselves rather than a reporter optimized for visibility. A potential improvement could use a Cre and flox design, with the enhancer driving Cre and the opsin and a soluble reporter on a floxed cassette. This would help to decouple brightness from enhancer strength and give both anatomical and functional measures in the same animal^9^.

### Developmental, viral and cell-intrinsic factors may all contribute to the PV bias

Several factors could contribute to the strong PV+ bias we observed. One factor could be that the *Dlx5/6* gene is broadly expressed throughout the developing GABAergic lineage. However it plays a particularly important role in the maturation and maintenance of PV-expressing interneurons after they become postmitotic^8,57,58^. In adult macaques where the developmental role of *Dlx5/6* in other interneuron subtypes has largely concluded, the *Dlx5/6* gene may remain more actively transcribed in PV+ neurons. Potentially explaining the strong PV-selectivity we and others observe^11^.

Another potential reason for this bias towards PV+ neurons might be that the AAV capsids themselves can carry intrinsic cell-type biases, independent of the enhancer that they package. AAV-PHP.eB has been shown via single-cell RNA sequencing to preferentially transduce PV+ and SST+ interneurons over VIP+ interneurons in mouse cortex, independent of any cell-type-specific promoter or enhancer^14,59^. While this may have contributed, it is unlikely to be the sole explanation. We observed strong preference for PV, modest transgene-expression in CB+ neurons and complete CR-avoidance using both AAV1 and AAV.PHP.eB capsids, two serotypes with distinct receptor-binding^60,61^ mechanisms and tropism profiles^62,63^. This pattern is not unique to our study either: similar PV-preference and CR-avoidance with the Dlx5/6 enhancer has been reported across macaque and rat cortex^11^ using PHP.eB, AAV1, and AAV9 serotypes, suggesting the bias is not attributable to our choice of capsids.

Yet another possibility is that cell types simply differ in how they handle AAVs^64^. PV+ neurons may be transduced more readily than CB+ neurons, and CB+ neurons more readily than CR+ neurons. There is precedent for such effects, with microglia resisting AAV transduction across both serotypes and promoters^64^. Relatedly, Dlx-based constructs may simply just drive stronger expression in PV+ and CB+ neurons. Thus, since opsin-reporter combinations are harder to express than fluorescent proteins alone, PV+ neurons may be easiest to detect as weakly labeled cells would be the first to fall below detection when the payload changes.

### Red-shifted opsins outperformed ChR2 for *in vivo* optotagging in the macaque

In our hands, the strongest *in vivo* optotagging responses came from ChrimsonR, followed by bReaChES, with ChR2 producing the weakest responses. We also observed that bReaChES was the most effective construct for *in vivo* suppression of neural activity. Differences in the robustness of *in vivo* optotagging can arise from several sources. Opsin-dependent variability in functional performance is well documented even within model organisms: a recent comparison of opsins in zebrafish found substantial differences in photocurrent amplitude, spiking induction, and high-frequency firing fidelity across opsins, even under matched expression conditions^65^. Another potential source is the intrinsic biophysical properties of the tissue itself. Blue light penetrates tissue less effectively than red light^66^, which could partly explain the weaker *in vivo* responses we observed with ChR2 relative to the red-shifted opsins ChrimsonR and bReaChES.

### Opsin and reporter are unlikely to be independently exchangeable parts

Yet another potential source for difference in optotagging performance is the combination of opsin and reporter used, which can impact trafficking and membrane localization^67,68^. We observed quantitative differences in expression patterns between construct payloads, with bReaChES/ChrimsonR-tdT exhibiting diffuse somatic fluorescence. Whereas ChR2-EYFP showed less somatic filling and more pronounced labeling of axonal processes, suggesting differences in intracellular trafficking or membrane localization. Increased functional opsin density at the soma could also be a potential explanation for why we observed that both red-shifted constructs, ChrimsonR-tdT and bReaChES-tdT, substantially outperformed ChR2-EYFP for driving action potentials. Other studies support the view that reporter and opsin interact in complex ways to affect function. In transgenic mice, ChR2(H134R)-EYFP was more photosensitive than ChR2(H134R)-tdT despite comparable membrane localization^67^. Whereas EYFP-tagged ChrimsonR showed higher photocurrent density than several other tagging strategies, including an untagged construct, in a separate *in vitro* study^68^. In both cases the tag altered function in ways that differed by opsin, suggesting that the consequence of a given reporter also depends on which opsin it is fused to. This reinforces our suggestion that the functional performance of these constructs depends not on opsin or reporter identity alone, but on the specific combination of the two.

### Response dynamics likely reflect opsin kinetics, not cell-type adaptation

Constructs also differed in the temporal dynamics of their optotagging responses. AiP12068 (ChrimsonR) showed a sharp initial peak in firing rate followed by a pronounced decay to a lower, sustained plateau, while AiP14858 (ChR2) showed a smaller, more gradually rising and sustained response, and ZE1001 (bReaChES) showed both increasing and decreasing firing rate populations with distinct kinetics (Adap. index, Fig. 3d-f). This pronounced adaptation might seem at odds with the canonical view of PV+ interneurons as minimally adapting, fast-spiking cells^69,70,71^. However, this expectation is largely derived from *in vitro* current-clamp experiments using fixed, step-like current injections, which differ substantially from *in vivo* optogenetic stimulation. Photocurrents from opsins desensitize under sustained illumination, with the degree and kinetics of this decay varying substantially across opsin variants^72^. Theoretical modeling of ChR2, ChrimsonR, and bReaChES shows that each displays a distinct peak-to-plateau photocurrent ratio and decay time course under constant light^73^. Because the resulting depolarization is not a fixed step input but instead reflects each opsin’s specific activation and desensitization kinetics, the firing dynamics we observe need not indicate adaptation intrinsic to the recorded cells.

### Cell health varies with payload

Finally, cell health also varied across constructs. Immunohistochemistry revealed that the majority of transgene-expressing cells were cleaved-caspase negative, indicating that transduction and expression were generally well tolerated. However, the ChR2-EYFP and bReaChES-tdT constructs elicited a greater proportion of cleaved-caspase+ cells than the ChrimsonR-tdT construct. This suggests that reporter^49,74,75^ and/or the protein fusion of opsin and reporter likely influence cell health in addition to their effects on trafficking and functional performance. We only performed this measure of cell health at one time, thus future studies could more systematically measure apoptosis signaling across time in order to better understand the progression and long term cell health consequences of these viral constructs.

### Viral vectors are akin to chemical formulas

Broadly, our results have three significant implications for the use of viral constructs in the study of inhibitory neuron function in primates. First, our results strongly suggest that capsid, enhancer, opsin, and reporter should not be treated as modular, independently swappable parts. Instead, viral constructs are better understood as something akin to chemical formulas, where the specific combination of components, not any single piece in isolation, determines functional, selectivity, and viability outcomes. Second, multimodal characterization as performed here is necessary for identifying the best performing tools for study of inhibitory neuron function. *In-vivo* recordings, immunohistochemistry or molecular characterization in isolation provide only pieces of the puzzle. Finally, our findings suggest that while there are currently abundant resources being dedicated into the identification of new enhancers, this may only be part of the solution towards cell-type specific construct development for the primate. Systematic, head-to-head comparison of capsid, opsin, and reporter combinations, as we and others have painstakingly pursued, are critical for moving the field forward.

### Summary and Conclusion

Taken together, our *in vivo* and *in vitro* data demonstrate the efficacy of three DLX5/6 enhancer-based viral constructs to genetically target and optogenetically modulate GABAergic neurons across macaque cortex. However, we found that these constructs do not lead to pan-inhibitory transgene expression. But rather preferentially express in select GABAergic subpopulations --- a finding consistent with other reports. As such, the promise of a single, perfectly pan-inhibitory viral construct for NHPs remains unrealized. Our findings thus emphasize the critical need to systematically characterize enhancer, capsid, and payload combinations using the multimodal approach described here.

This characterization approach has several benefits. First, each characterization experiment will generate ground truth *in vivo* optotagging data linking waveform and firing properties to known cell identity. Second, such data combined across experiments can be used to train cell-type classifiers capable of identifying multiple inhibitory subtypes simultaneously, even for *in vivo* datasets that lack ground truth. Third, validated tools can allow for GABAergic circuit-specific functional mapping and modulation *in vitro* for mechanistic characterization of GABAergic synaptic signaling and pharmacology in primates. Iterating these efforts of tool development, generating ground truth mechanistic data, and circuit-specific functional mapping will build upon each other and move the field closer to understanding how the diverse inhibitory cell types of the primate brain sculpt circuit dynamics underlying cognition and behavior.

## 4. Methods

### 4.1. Subjects

For the experiments reported here, we used 7 adult rhesus macaque monkeys (*Macaca mulatta*; 5 females and 2 males, 6 - 21 years old Fig. S3)). We identified monkeys with complete health records from national primate research facilities and from private vendors. All animals used here are part of our larger studies of normal brain aging and tract tracing. We housed the animals individually in the Animal Science Center (ASC) at Boston University Medical Campus under a 12-hour light/dark cycle. All of our procedures were approved by the Boston University Institutional Animal Care and Use Committee (2018-00053) and conducted in strict accordance with the guidelines established by the NIH Guide for the Care and Use of Laboratory Animals.

### 4.2. Viral injections in macaques

We injected viral constructs into targeted brain areas, using MRI, surgical, and tissue processing procedures described previously^76–78^. Injection sites were based on stereotaxic coordinates calculated from MRI scans taken prior to surgery, using the midline and ear canals as reference points, as described^79^. For MRI, monkeys were sedated with a mixture of ketamine (10 mg/kg) and valium (0.5-1.0 mg/kg, to effect), which was followed with additional doses as needed. For injection site planning, a structural MRI was acquired using a T1-weighted 3D-turbo field echo (TFE) sequence, fully optimized to provide high signal to noise ratio (SNR) and high gray-white matter contrast at high resolution with the following parameters: Repetition time (TR)/Echo time (TE) = 7 ms/3 ms, flip angle = 8°, number of excitations/averages (NEX) = 8, voxel size = 0.7 x 0.7 x 0.7 mm^3^, sagittal plane acquisition. Additional scan sequences were obtained from each monkey as part of parallel studies.

For each surgery, monkeys were sedated with ketamine (10mg/kg, to effect) and anesthetized to the surgical plane with isoflurane (induction at 2-3%, to effect). For each animal, two surgeries (at least 3 weeks apart) were made to perform a craniotomy and expose one hemisphere per surgery (one hemisphere was for *in vivo* recordings and the other for *in vitro*, Fig. S3). For monkeys W and M there was only one surgery on one hemisphere where *in vivo* and *in vitro* recordings were performed in the same tissue. For each surgery, monkeys were intubated and placed in a stereotaxic head holder. Surgery was performed under sterile conditions, with vitals (blood-oxygen saturation, heart rate, respiratory rate, and temperature) continuously monitored. Craniotomies were made per hemisphere spanning the caudal-lateral prefrontal cortex (LPFC area 46 and 8), ventral and dorsal premotor and primary motor cortices (PMd, PMv and M1) and primary somatosensory cortex (S1). The dura was reflected to provide access to the underlying brain tissue. 10 µl Hamilton (Hamilton, Reno, NV, USA) syringes preloaded with viral construct were carefully stereotaxically lowered 2-3 mm into the target brain area (Fig. S2a). Viral constructs were sourced from viral cores at the Allen Institute (Seattle, WA, USA), University of Texas at Austin (author BZ’s lab, Austin, TX, USA) or from PackGene (Houston, TX, USA). 2.2-3.2 µl of viral construct were injected per penetration to the target brain area at a rate of 500 nL per minute using a syringe pump (UMP3T-1 with SMARTouch Controller, World Precision Instruments, Sarasota, FL, USA). Following the injection of the viral construct, the Hamilton syringe remained *in situ* for 5 minutes to allow the viral construct to diffuse through the tissue and prevent upward suction of the viral construct upon retraction of the needle. 2-3 penetrations and injections were performed for each target brain area to inject a total of 6-9 µl per brain area. Following the injections, the dura was repositioned and the skull fragment was sutured back into the rest of the skull. Post-injection, all monkeys made a full recovery while the viral constructs incubated in the target brain regions for 44-156 days (Fig. S3).

### 4.3. Anesthetized in vivo recordings in macaques

Monkeys were anesthetized via a ketamine injection (10mg/kg, to effect). Monkeys were kept warm with heating blankets and intubated to control breathing. Anesthesia was maintained with continuous release propofol (2.5-5.0 mg/kg, IV). Vitals (blood-oxygen saturation, heart rate, and breathing rate) were continuously monitored throughout the experiment to ensure the animal’s safety and to ensure the animal was not experiencing any distress. Craniotomies were performed to remove portions of the skull to expose the underlying cortical structures of interest and the dura reflected to provide access to the underlying brain tissue.

All optotagging experiments, except one, reported here were performed with a 32-channel Plexon (Dallas, TX, USA) S-probe (100 µm vertical spacing between channels; fitted with an optical port between channels 16/17, Fig. S2b). The Plexon S-probe’s optic fiber has an internal diameter of 46 µm, outer diameter 50 µm, and a .66 numerical aperture. Thus the programmed power of the laser (10-330 mW) was typically reduced 90-95% upon exiting the probe, leading to effective stimulation powers of 0.5-5 mW. One experiment (Monkey S, 13 units) was performed with a 128-channel Diagnostics Biochips (Glen Burnie, MD, USA) DA128-2 probe (dual column channel layout with 43.3 µm horizontal spacing, 50 µm same-column vertical spacing, and 25 µm alternate-column vertical spacing between channels; fitted with an optical port past the first 3rd of channels of the most superficial channels). The DA128-2 probe’s optic fiber has an internal diameter of 200 µm, outer diameter of 245 µm, and a .37 numerical aperture.

Closed-path laser stimulation was achieved with Thorlabs (Thorlabs, Newton, NJ, USA) fiber optic patch cables connected from the laser head and mounted to the probe via a ferrule at the top of the probe. Path cables were 1 m in length, with a 200 µm core diameter, .22 numerical aperture and a 2.5 mm FC/PC connection to the laser bulkhead and 1.25/2.5 mm ferrule attachment to the Plexon S-probes fitted together with a mating sleeve (Thorlabs, Newton, NJ, USA). Three lasers were used throughout these experiments: either a gem 473 nm, 250 mW or gem 561 nm, 250 mW (Novanta, Bedford, MA, USA), or a LuxX+ 638 nm, 200 mW (Omnicron, Rodgau, DE).

All lasers were operated with serial commands to turn them on/off. Generally, lasers were kept on during the duration of a stimulation block and laser pulses were delivered via shutters. The analog signal to control laser pulsing was generated with a Blackrock Microsystems Cerebus system (Blackrock Neurotech, Salt Lake City, UT, USA). The analog signal was split to open/close the shutter and to record the timing of the analog signal via the Open Ephys I/O board^80^. gem lasers were generally operated using a 1x1 fiber-fiber optical switch (model FFLS, Agiltron, Woburn, MA, USA) to precisely control the timing of laser onset (∼15 ms delay) and offset (∼12 ms delay).

Several other systems of laser pulse control were used with the gem laser systems prior to settling on the Agiltron shutter solution. In initial experiments (W, M and T), serial commands were used to control gem laser pulsing via turning the laser on/off. However, the delay between the timing of the serial command and when the laser actually turned on/off was variable and therefore unreliable to use to align trials. Thus, one subject’s (W) recordings were aligned to stimulus artifact onset and offset. For M and T a 99:1 fiber optic patch cable splitter (Thorlabs) was used with 1% of power going to a photo diode to exactly record the laser on period. For monkey B, a custom interlock with a laser onset and offset delay of ∼28 ms and ∼12 ms respectively was used. All laser onset and offset times were appropriately accounted for and properly aligned to the neural signal. These timings were additionally confirmed with the onset and offset of photovoltaic artifact.

Probes were carefully lowered 3 - 5 mm into the targeted cortical brain area (e.g., PMdc/M1) such that all recording channels would be in neocortex. Electrophysiological recordings were made using Open Ephys hardware^80^ and the Open Ephys GUI^80^. Raw voltage and band-pass Butterworth filtered (300-6,000 Hz) spiking activity were sampled and saved at 30 kHz. As voltage recordings occurred in an operating room, care was taken to improve the quality of the recording (e.g., adjusting the ground screw) before recording baseline brain activity for 30 seconds - 5 minutes. After baseline recording, stimulation recordings were performed where probe depth, stimulation power, and stimulation protocol were varied to optimize optotagging yield. Once optimal optotagging was determined, the most common stimulation protocol involved 50 bursts of 5, 50 ms square-wave pulses with a 250 ms inter-pulse interval, and 1.5 s inter-burst interval for a total of 250 pulses (∼2.5 minutes). Several stimulation procedures were performed for each penetration in a brain area.

### 4.4. Perfusion and preparation of acute slices

Immediately after surgical recordings, harvesting of acute brain slices for *in vitro* recordings followed by whole brain fixation was performed using our standard two-stage Krebs-Paraformaldehyde perfusion method, as described^81^. While under surgical level of anesthesia and with the monkeys’ head fixed in a stereotactic head holder, the craniotomy was expanded to provide access to the left hemisphere for removal of fresh biopsy samples, as described previously^81^. Following administration of a final dose of propofol (2.5 - 5.0 mg/kg), the monkeys were then perfused through the heart first with ice-cold Krebs–Henseleit buffer (concentrations, in mM: 6.4 Na_2_HPO_4_, 1.4 NaH_2_PO_4_, 137 NaCl, 2.7 KCl, 5 Glucose, 0.3 CaCl_2_, 1 MgCl_2_; pH 7.4, Sigma-Aldrich, St. Louis, MO, USA). With Krebs buffer flowing, one to two blocks of tissue (10 mm^3^) were taken from the PMdc/M1 and S1. The blocks were removed from the hemisphere designated for *in vitro* recordings and sectioned into 300 µm thick coronal slices in ice-cold oxygenated (aerated with 95% O_2_, 5% CO_2_) NMDG solution, with a vibrating microtome as modified from our previous protocols^82^ and Ting et al, 2018^83^. After slicing an entire block (∼15 min in ice cold NMDG) slices were then transferred from ice-cold to 33 ^°^C oxygenated NMDG solution for 10 min, then immediately placed into room temperature, oxygenated Ringer’s solution (concentrations, in mM: 26 NaHCO_3_, 124 NaCl, 2 KCl, 3 KH_2_PO_4_, 10 glucose, 1.3 MgCl_2_; pH 7.4; Sigma Aldrich). After blocks were dissected, the remainder of the left hemisphere was quickly immersion-fixed in freshly depolymerized 4% paraformaldehyde (in 0.1 M phosphate buffer [PB], with 0.9% NaCl, pH 7.4) at room temperature for 24 hours, then transferred to fresh 4% PFA and kept at 4 ^°^C until use. Blocks containing the injection sites from the left hemisphere were dissected then sectioned at 100 µm in the vibratome for injection site validation.

Once fresh tissue harvesting was concluded, perfusate was switched to freshly depolymerized 4% paraformaldehyde (EM grade; Ladd Research Industries, Williston, VT, USA) in 0.1 M PB with 0.9% NaCl (pH 7.4, at 37 ^°^C) to fix the intact hemisphere designated for *in vitro* recordings for injection and recording site validation, serial sectioning for immunohistochemistry and 3D mapping of labeled cells. The hemisphere was removed from the skull and post-fixed for 24 hours at 4 ^°^C, in 4% paraformaldehyde. The hemisphere was blocked at the coronal plane, cryoprotected in graded solutions of glycerol (10 – 30%), frozen in -70 ^°^C isopentane and sectioned on a freezing microtome in the coronal plane at 60 µm, as described^78^. All fixed vibratome slices and thin tissue sections were stored in antifreeze solution (30% ethylene glycol, 30% glycerol, in 0.05 M PB, pH 7.4) at -20 ^°^C until use.

### 4.5. Whole-cell patch clamp electrophysiology on acute in vitro slices

After at least a 1-hour equilibration period, individual acute slices were placed into submersion-type recording chambers (Harvard Apparatus, Holliston, MA, USA) mounted on the stages of Nikon E600 infrared differential interference contrast (IR-DIC) microscopes (Micro Video Instruments, Avon, MA, USA). Slices were continuously superfused with room temperature, oxygenated Ringer’s solution at a rate of 2-2.5 mL/min. All physiological recordings were performed at room temperature (∼26 ^°^C), which improves the viability and duration of recordings from monkey slices. Standard tight-seal, whole-cell patch-clamp recordings with simultaneous biocytin filling were obtained from unlabeled L3 and L5 pyramidal neurons and construct-labeled interneurons from S1, PFC area 46, MCC area 24c as described previously^81,84–86^. Cells were visualized first under epifluorescence illumination to identify construct-labeled neurons. Once a labeled cell was identified, optics were switched to IR-DIC optics for approaching with patch electrodes fabricated on a horizontal Flaming and Brown micropipette puller (Model P-87, Sutter Instruments, Novato, CA, USA). Potassium methanesulfonate-based internal solution (concentrations, in mM: 122 KCH_3_SO_3_, 2 MgCl_2_, 5 EGTA, 10 NaHEPES, with 0.5% biocytin, pH 7.4; Sigma-Aldrich) was used to fill patch electrodes with resistances of 3-6 MΩ in the external Ringer’s solution. Data were acquired using EPC-9 or EPC-10 patch-clamp amplifiers (HEKA Elektronik, Lambrecht, Germany), using PatchMaster acquisition software (HEKA Elektronik). The series resistance ranged from 10-15 MΩ and was not compensated. Series and access resistance were both frequently monitored during the 15-20 min recording period for each cell. If there was a >10% change in series or access resistance, the cell was discarded. Bessel filter frequency was 10 kHz and sampling frequency was at 7 kHz for voltage clamp and 12 kHz for current clamp recordings.

#### Optogenetic stimulation to assess optotagging

We performed whole-cell patch-clamp recordings from optotagged inhibitory neurons, identified via fluorescent labeling with tdT- or EYFP-reporter protein. The standard pipeline using a series of current injections was used to measure intrinsic and firing properties from each cell. First, a repeated short 5-20 ms optical pulse was delivered at ∼0.65-3 mW with either a collimated LED light source (cool LED ultra) (CoolLED Ltd., Andover, NY, USA) or fiber optic cannula (Thorlabs, Newton, NJ, USA) coupled laser source (same as for *in vivo* recordings, see *4.3*) to test if the cell was optotagged (fired an action potential time-locked to the light pulse). If a recorded cell was optotagged, the following protocols were run to measure voltage responses: First, a series of 50 ms light pulses (interstimulus interval (ISI) 5 s) with increasing power from 0.65 mW to 3 mW LED power were run to create an input-output curve and identify the minimum stimulus to elicit an action potential. Then at minimum light power a repeating 10 series of light pulses at 50 ms, 100 ms and 200 ms duration and 5 s ISI were run. Then we ran a protocol to mimic the *in vivo* stimulus: 200 ms duration light pulses with 100 ms ISIs, repeated 4 times per sweep.

#### Optogenetic stimulation of inhibitory fibers

To assess optically-evoked inhibitory synaptic transmission, we recorded from unlabeled pyramidal neurons surrounded by tdT- or EYFP-labeled fibers. We first measured intrinsic and firing properties, followed by voltage clamp measurements of spontaneous inhibitory synaptic currents. We then determined if the pyramidal neuron was synaptically coupled to inhibitory terminals by holding the cell under voltage clamp at -40 mV and delivering a 5-20 ms light pulse at ∼0.65-3 mW light power. A cell was considered “coupled”, if an inhibitory postsynaptic current (IPSC) was elicited time-locked to the light stimulation (thus optically-evoked IPSC (oIPSC)). Once coupling was determined, the following protocols were run under voltage clamp mode to measure oIPSCs: First, a series of 5 ms light pulses (ISI 5 s) with increasing power from 1 to 10% LED power (∼0.65-3 mW; 470 nm, 565 nm, 590 nm, coolLED pE-300ultra) or 2 to 25% laser power [2 sweeps per power; gem 473 nm, 250 mW, gem 561 nm, 250 mW (Novanta), and a LuxX+ 638 nm, 200 mW (Omnicron)] were run to create an input-output curve and identify the minimum stimulus to elicit oIPSCs (Fig. 6d). Then, at this minimum light power, a repeating 10 series of light pulses at 5 ms (5 s ISI) was run to measure the average oIPSC waveform. Then paired-pulse (2 pulses at 200 ms ISI) and multi-pulse (4 pulses at 200 ms ISI) protocols were run to measure short-term presynaptic plasticity dynamics of the oIPSCs (facilitation or depression).

#### Analysis pipeline for *in vitro* electrophysiological data

Analyses of passive membrane and action potential firing properties were conducted using custom MATLAB scripts, as described in our previous work^87^. Specifically, resting potential (Vr) membrane time constant (tau), input resistance (Rn), voltage sag potential amplitude (Vsag), and action potential waveform properties (amplitude, rise fall, duration at half max amplitude) were measured for each recorded pyramidal neuron and optotagged inhibitory interneuron. To further characterize the recorded cells, the frequency, amplitude and kinetics (rise, decay and duration) of spontaneous postsynaptic excitatory/inhibitory currents (sEPSCs and sIPSCs respectively) were measured using Minianalysis software, as previously described^87^. For oIPSCs recorded in synaptically coupled pyramidal neurons, peak amplitude, kinetics (rise time, decay time, duration), and latency were measured using custom MATLAB scripts. The paired-pulse ratio from each cell was calculated as the ratio of peak amplitude of the oIPSC response of pulse 2 divided by peak amplitude of oIPSC response to pulse 1 (peak 2/peak 1).

#### Slice processing to visualize recorded cells

Following whole-cell patch clamp recordings, slices were fixed in 4% paraformaldehyde in 0.1 M phosphate-buffered saline (PBS; pH=7.4) for 48h at 4 °C. Subsequently slices were incubated for 2 h in 1% Triton X (in 0.1 M PBS), followed by 48 h in streptavidin-Alexa 750 (diluted 1:500 in 0.1 M PBS; Thermo Fisher). Finally, slices were washed 3 x 10m in 0.1 M PBS then stored at -20 °C in anti-freeze solution (comprised of 30% ethylene glycol (Sigma), 30% glycerin (Sigma), 40% PB at a final concentration of 0.05 M, ph 7.4) until confocal imaging as described below.

### 4.6. Injection site validation

Gross validation of injection sites were performed at the time of tissue harvest and fixation. For each case, the intact lateral and medial surface of each whole hemisphere was photographed by combining the Xite fluorescence LED (NIGHTSEA, Morgantown, PA) and the standard camera on an iPad. The coordinates and details of each injection site are mapped onto a schematic brain surface and summarized in (Fig S3).

Further histological validation was performed for a subset of cases as follows: For the cases used for *in vitro* recordings (Mky M, A, and C), slices containing the injection sites were fixed and examined under an epifluorescence microscope for validation of the spread of the label. For the fixed intact hemisphere used for *in vivo* recordings, a subset of cases (Mky B, C, A, and S) were used for serial section analyses of label and immunohistochemical validation experiments. A series of coronal sections cut on a freezing microtome (60 µm, 300µm apart) were examined to identify the sections with the injections. These sections were photographed under epifluorescence and then medio-lateral span was measured using area measurements in FIJI, and the rostro-caudal span was calculated based on the total number of sections spanning the injection site, and the spacing between sections (Fig. S4a). Two of the brains (Mky B and S) were then used for visualization of injection sites onto the 3D whole brain surface using 3D Slicer^88^ (see Fig S4b-g). For this analysis, MRI T1 scans of the brains (at 0.7 x 0.7 x 0.7 voxel, see methods above) were imported into 3D slicer and images of coronal sections with injection sites were aligned onto corresponding MRI slices. After transformation and aligning of labeled histological sections onto each corresponding reference MRI slice, injection sites were traced in each section and rendered in 3D reconstruction for visualization (Fig. S4c, f).

### 4.7 Immunohistochemistry and quantification of optotagged cells

For a subset of cases (Mky B, S, A and C, see Fig. S4), quadruple-label fluorescent immunohistochemistry on free floating 60 µm sections was performed to examine the density and distribution of AiP12068+, AiP14858+, and ZE1001+ cells colabeled for the calcium-binding proteins calretinin (CR), calbindin (CB), parvalbumin (PV) and/or the vesicular GABA transporter (VGAT). In a second quadruple immunohistochemistry set, tissue sections containing AiP12068+, AiP14858+, or ZE1001+ cells were immunostained for markers of gliosis (Iba1 for microglia), GFAP (for astrocytes) or cell death (cleaved caspase) as well as for GAD67 (pan-interneuronal marker). For every animal and cortical region, three sections spaced 300 μm apart were processed. Tissue sections were initially rinsed in 0.01 M PBS (three 10-min washes at 20 °C) and then incubated for 1 h at 20 °C in 50 mM glycine in 0.01 M PBS. Antigen retrieval involved placing sections in 10 mM sodium citrate buffer (pH ∼8.5) in a 60-70 °C water bath for 20 min, followed by a 30-min cool-down on a shaker at approximately 20 °C. After another series of PBS rinses (3 × 10 min), sections were pre-blocked for 1 h at 20 °C in a solution of 0.01 M PBS containing 5% bovine serum albumin, 5% normal goat serum, and 0.2% Triton X-100.

Primary and secondary antibodies used in this study have been validated by their manufacturers, are widely cited, and are listed in Table 1. Primary antibodies were diluted in a buffer of 0.1 M PB supplemented with 0.2% acetylated BSA, 1% normal goat serum, and 0.1% Triton X-100. Two sets of antibody combinations were used. In the first set, tissue sections were incubated simultaneously in rabbit anti-CR (Swant, Kehrsatz, Switzerland) 1:500; goat anti-CB (Thermo Fisher Scientific, Waltham, MA, USA) 1:100; guinea pig anti-VGAT (Synaptic Systems, Goettinggen, Germany) 1:400 and mouse anti-PV (Swant) 1:1000. The second set included guinea pig anti-IBA1 (Synaptic Systems) 1:500; goat anti-GAD67 (R&D Systems, Minneapolis, Minnesota) 1:100; mouse anti-GFAP (Synaptic Systems) 1:500 and a rabbit anti-cleaved caspase 3 direct conjugate (Cell Signaling, Danvers, MA, USA) 1:100. For both experimental sets, sections were exposed to two 10-min microwave cycles at 150 W and 40 °C using a PELCO Biowave (Ted Pella, Inc., Redding, CA), followed by a 72 h incubation at 4 °C with continuous moderate agitation on a shaker platform.

Following incubation in primary antibody solution, sections were washed in PBS (3 × 10 min) and incubated in their respective secondary antibody mixtures using the same microwave-assisted protocol (2 × 10-min cycles, 150 W, 40 °C), then incubated on a shaker at 4 °C for 24 h. Secondary antibodies (Table 1) for Set 1 (interneuron markers) included: donkey anti-goat F(ab’)2 405 (1:200); donkey anti-rabbit F(ab’)2 488 (1:200); donkey anti-guinea pig F(ab’)2 647 (1:200); and donkey anti-mouse F(ab’)2 750 (1:200). For Set 2 (glia and cleaved caspase markers) included: donkey anti-guinea pig F(ab’)2 DyLight 405 (1:200); donkey anti-goat IgG 488 (1:200); and donkey anti-mouse IgG 750 (1:200). After final rinses in 0.1 M PB (3 × 10 min at 4 °C), sections were mounted on slides and coverslipped with ProLong Gold Antifade (Thermo Fisher Scientific).

#### Confocal laser scanning microscopy

Tissue was imaged with an Olympus Fluoview FV3000 confocal microscope (Olympus, Tokyo, Japan). To quantify densities of single- and double- or triple-labeled tdT+ or EYFP+ neurons, z-stack images were acquired using a 20x objective (air, 0.75 NA). A 1272 (2 x 636)-μm-wide cortical column extending from the pial surface to the white matter was captured for each region and section containing tdT+ or EYFP+ neurons (Fig. 2). Image stacks were collected with a voxel size of 0.6215 x 0.6215 x 0.3 μm. Five channels were acquired independently using lasers at 405 nm, 488 nm, 561 nm, 647 nm and 750 nm. Deconvolution of image stacks was performed using AutoQuant (as in Gilman et al., 2017; Tsolias & Medalla, 2022^82,89^). For each subject, three sections were imaged for each region of interest (ROI). To further assess expression level of reporter gene and of cleaved caspase within individual cells, we imaged a subset of fields (2 per section) at a higher resolution at 40x, oil immersion, 1.4 NA (0.155 x 0.155 x 0.3 µm voxel size).

#### Quantification of single- and colabeled-tdT+ or EYFP+ neurons

Each coronal section, a total of 2 columnar ROIs (1272 µm wide) spanning the entire cortical depth for each area, were imaged at 20x (0.6215 x 0.6215 x 0.3 μm) at the confocal microscope. For each case, a total of 6 columnar ROIs across 3 coronal sections (2 ROIs per section) were imaged and quantified for reporter+ cell bodies. Neuron counts spanning the full cortical depth of each z-stack montage were performed in QuPath (Version: 0.4.2). In total, 403-711 tdT+ (AiP12068 Mky B PMdc/M1 n = 711, Mky S n = 619, Mky A n = 407; ZE1001 Mky C n = 403) and 120 EYFP+ (Mky B S1) neuronal cell bodies were counted per case. All tdT+ or EYFP+ neurons with nonpyramidal morphology were exhaustively identified within each columnar ROI and assessed for the co-expression of calcium binding proteins. For this analysis of cell numbers and proportions, the maximum z projection image with 51 optical slices (15µm thickness) from each confocal stack ROI was imported into QuPath. The tdT+ cell bodies were detected using the automated cell detection function (with manual editing) in QuPath and were categorized based on co-localization with one or two of the calcium binding proteins. Individual EYFP+ neurons were identified and demarcated manually based on a threshold of signal intensity (automatic cell detection protocols are not able to detect the low levels of EYFP in somata against the strong neuropil labeling). Reporter+ cells were categorized as either reporter only (single labeled), or reporter co-labeled with one [CR+, CB+ or PV+ (dual-labeled)] or a combination of two [CR+/CB+, CR+/PV+, CB+/PV+ (triple-labeled)] calcium binding proteins. The same analysis was conducted on a subset of cases (AiP12068 Mky B PMdc/M1, Mky S S1, ZE1001 Mky C S1 tdT, AiP14858 Mky B S1 EYFP) to assess reporter+ neurons co-labeled with GAD67 and Cleaved Caspase 3 (ClvCas3). However, due to the punctate nature of GAD67 and ClvCas3 labels, a second set of higher resolution images at 40x (0.155 x 0.155 x 0.3 µm voxel size) were obtained for analyses. For each coronal section, 3-4 ROIs in superficial and 3-4 ROIs in deep layers within each labeled region were imaged. A total of 20-24 40x fields across 3 sections were quantified per case. The z-maximum projection image with 21 optical slices (6.3 µm in z thickness) from each confocal stack ROI was imported into QuPath for quantification of reporter+ somata co-localized with GAD67+ or ClvCas3, as described above.

In this study we define completeness as the percentage of a population that is colabeled with a construct reporter. For example, what percentage of all GABAergic cells were also labeled by a viral construct? Specificity is defined as the percentage of all reporter+ cells that were colabeled with a population marker. For example, what percentage of all ChrimsonR-tdT+ neurons were also labeled by a cell population marker. Specificity and completeness are formalized below:

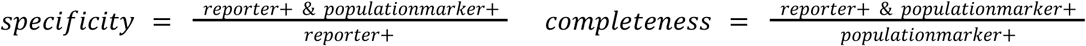

In this second immunohistochemistry set with reporter, GAD67, cleaved caspase and Iba1 and GFAP, we performed a second analysis to assess expression intensity of reporter proteins and how it relates to inflammatory glial markers, Iba1 and GFAP. For this glial analysis, 20x columnar ROI imaging was performed as with the set 1 immunohistochemistry (with CB, PV, and CR). The glial markers, Iba1 and GFAP, were quantified using our established particle analyses optical density method, as described^87^. Thresholds for each channel were set using semi-automated methods in FIJI, and the particle analysis plug-in in FIJI was employed to quantify percent area labeled by Iba1, GFAP and tdT. This optical label density measure of tdT was then correlated with density of Iba1 or GFAP label to assess relationship between inflammatory glial markers and extent of viral transduction.

To examine laminar distributions, we drew contours marking the pial and white matter boundaries for each section. We then measured and expressed the distance of each labeled neuron from the pia as a percentage of total cortical depth. To understand the depth distribution for each neuronal population, we generated histograms of normalized neuron counts (with bins of 100-µm increments of distance from the pial surface).

### 4.8. In-vivo data processing pipeline and spike sorting

We loaded voltage recordings into MATLAB (The MathWorks, Inc., Natick, MA, USA) using the open-ephys-matlab-tools^90^ toolbox. We wrote custom MATLAB scripts to preprocess and visualize recorded voltage data. We first band-pass filtered the raw voltage (Butterworth, 300-6,000 Hz) and used common average referencing to remove noise. If the recording probe remained at the same depth between adjacent recordings, we combined recordings to increase the number of trials and therefore improve our ability to find responsive units to laser stimulation.

After preprocessing the voltage data, we used Kilosort4 for automated spike sorting^34^. We then used manual curation to further refine acquired clusters. We excluded obvious noise clusters or non-neural waveforms. We only included and analyzed units that were modulated in relation to laser stimulation (i.e., optotagged or suppressed). We did not include units in our analyses that were spontaneously active during the recording but had no discernible response to laser onset. To determine if a unit was optotagged, we performed the stimulus associated latency test (SALT)^3^. Generally, we found that SALT performed well, appropriately identifying optotagged cells. However, we performed additional manual curation if a unit ‘responded’ particularly weakly. To determine if a unit was suppressed, we used a similar method to SALT described as the “*Detection of light-induced inhibition*” as described previously^3^. For all clusters, we used manual inspection to remove any noise, and aligned waveforms to a common trough. If we were unable to align the waveforms, we did not include the cluster for further analysis. We found that photovoltaic artifacts were minimal and typically only appeared at the onset and offset of laser timings. We found that Kilosort4 did not typically identify photovoltaic artifacts as clusters. However, if it did, we excluded these artifact clusters as noise.

### 4.9. Peristimulus time histograms, response latency and waveform preprocessing

To calculate peristimulus time histograms, we aligned spike times to laser onset for each pulse (i.e., in Fig. 3a, b, c). We then convolved these full sampling resolution (30000 samples/s) spike trains with a normalized Gaussian kernel (σ= 0.3 ms; truncated at ± 3σ) and estimated the instantaneous firing rate (spikes/s). Finally, we averaged the firing rates across all laser pulses to estimate the response to optical stimulation. We then averaged peristimulus time histograms from all neurons per construct (as in Figure 3d, e, f). To estimate the response latency per construct, we first calculated the first time that a responsive unit’s peristimulus time histogram reached the half-peak response after the onset of the laser. We then estimated the median latency across units. Adaptation index was calculated as the ratio of late (i.e., 40-50 ms) to early (i.e., 20-30 ms) median firing rates during the stimulation period per unit. To calculate waveform correlation we averaged all the spontaneous waveforms and separately averaged evoked waveforms of an optotagged unit. Then we calculated the Pearson correlation of the average spontaneous and evoked waveforms.

To prepare *in vivo* waveforms for analysis in WaveMAP^25^, a waveform shape clustering algorithm, we performed a number of preprocessing steps (Fig. 4a). First, we spike-sorted and manually curated unit waveforms. We included units whose firing rate either increased or decreased in response to the laser, as described above. One additional unit from monkey M did not pass the SALT test (∼17 ms response time post laser onset) but was included in the WaveMAP analysis. The unit was clearly responsive after the onset of the laser and we believe its delayed response was due to a technical issue (i.e., damaged fiber optic ferrule on the V-probe) rather than a biological delay. We also included 3 units from monkey W that passed the SALT test but showed a paradoxical effect of suppressing their activity 3/4s of the way through the laser stimulation period. We also removed any noise waveforms from responsive clusters. Finally, we averaged waveforms per unit, interpolated them to an effective sampling rate of 120,000 samples/s, normalized their amplitude and polarity, and centered them at their trough, keeping 1 ms before and 2 ms after.

We pursued similar steps to prepare *in vitro* waveforms for analysis in WaveMAP^25^. We included *in vitro* cell waveforms from two sources: 1) when the cell expressed a pan-inhibitory transgene from one of the three constructs and was either synaptically-coupled to an optogenetically-tagged fiber (n = 9) or optotagged (n = 4) or 2) when a biocytin-filled cell from a previous recording was visually identified as having either a pyramidal (excitatory) or interneuron (inhibitory) morphology (n = 37). Selected neurons had a train of action potentials (> 1 action potential) in response to a 2 s current injection. We chose the third action potential in the train as the representative waveform of that cell (in a few cases there were only two action potentials and the second action potential was used). If there were multiple candidate spike waveforms to be chosen from due to multiple sweeps, the largest amplitude waveform was selected. We again interpolated waveforms to an effective sampling rate of 120,000 samples/s, amplitude normalized them to 1, and centered them at their peak with 3 ms before and 6 ms after.

## Acknowledgements

We thank the Viral Technology Core team at the Allen Institute for AAV packaging services. B.P.L. and J.T.T. were supported by NIH BRAIN Initiative (UG3/UH3 MH120095). The goal of this grant is to support development and dissemination of cell type specific AAV vectors for cross species functional studies. The following authors were all supported by the National Institute of Neurological Disorders and Stroke: CC (R00 NS092972, R01 NS121409, R01 NS122969, R21 NS135361), JIL (R01-NS122969), EKL (F31 NS131018), and RJR (R01 NS125307). CC was additionally supported by the Whitehall Foundation (2019-12-77), the Brain and Behavior Research Foundation (27923), and the National Science Foundation (CAREER INSF IOS-2541566). JIL was additionally supported by the National Institute on Aging (R01 AG092516, R21 AG084041). MM was supported by the National Institute on Mental Health (R01 MH116008) and MM and TLM were supported by the National Institute on Aging (R01 AG083206).

## References

1. Krimer, L. S. et al. Cluster analysis-based physiological classification and morphological properties of inhibitory neurons in layers 2-3 of monkey dorsolateral prefrontal cortex. J Neurophysiol 94, 3009–3022 (2005).

2. Lewis, D. A., Hashimoto, T. & Volk, D. W. Cortical inhibitory neurons and schizophrenia. Nat Rev Neurosci 6, 312–324 (2005).

3. Kvitsiani, D. et al. Distinct behavioural and network correlates of two interneuron types in prefrontal cortex. Nature 498, 363–366 (2013).

4. Pi, H.-J. et al. Cortical interneurons that specialize in disinhibitory control. Nature 503, 521–524 (2013).

5. Courtin, J. et al. Prefrontal parvalbumin interneurons shape neuronal activity to drive fear expression. Nature 505, 92–96 (2014).

6. Estebanez, L., Hoffmann, D., Voigt, B. C. & Poulet, J. F. A. Parvalbumin-Expressing GABAergic Neurons in Primary Motor Cortex Signal Reaching. Cell Rep 20, 308–318 (2017).

7. De, A., El-Shamayleh, Y. & Horwitz, G. D. Fast and reversible neural inactivation in macaque cortex by optogenetic stimulation of GABAergic neurons. Elife 9, (2020).

8. Dimidschstein, J. et al. A viral strategy for targeting and manipulating interneurons across vertebrate species. Nat Neurosci 19, 1743–1749 (2016).

9. Mehta, P. et al. Functional Access to Neuron Subclasses in Rodent and Primate Forebrain. Cell Rep 26, 2818–2832.e8 (2019).

10. Mich, J. K. et al. Functional enhancer elements drive subclass-selective expression from mouse to primate neocortex. Cell Rep 34, 108754 (2021).

11. Yazdan-Shahmorad, P., Gibson, S., Lee, J. C. & Horwitz, G. D. Preferential transduction of parvalbumin-expressing cortical neurons by AAV-mDLX5/6 vectors. Front Neurosci 17, 1269025 (2023).

12. Li, L. et al. Identification and application of cell-type-specific enhancers for the macaque brain. Cell 188, 4471–4473 (2025).

13. Federer, F. et al. Laminar specificity and coverage of viral-mediated gene expression restricted to GABAergic interneurons and their parvalbumin subclass in marmoset primary visual cortex. Elife 13, (2024).

14. Furlanis, E. et al. An enhancer-AAV toolbox to target and manipulate distinct interneuron subtypes. Neuron 113, 1525–1547.e15 (2025).

15. Vafaei, A. et al. Laminar-specific control of response gain and orientation-tuning by parvalbumin-expressing inhibitory interneurons in primate visual cortex. Res Sq (2026) doi:10.21203/rs.3.rs-9117440/v1.

16. Tremblay, S. et al. An Open Resource for Non-human Primate Optogenetics. Neuron 108, 1075–1090.e6 (2020).

17. Deisseroth, K. Optogenetics: 10 years of microbial opsins in neuroscience. Nat Neurosci 18, 1213–1225 (2015).

18. Su, C. & El-Shamayleh, Y. Optotagging in primate cortex uncovers excitatory and inhibitory neuron specializations in form vision. Nature Communications 1–20 (2026).

19. Bliss-Moreau, E., Costa, V. D. & Baxter, M. G. A pragmatic reevaluation of the efficacy of nonhuman primate optogenetics for psychiatry. Oxf Open Neurosci 1, kvac006 (2022).

20. Li, N. et al. Spatiotemporal constraints on optogenetic inactivation in cortical circuits. Elife 8, (2019).

21. Fishell, G. & Kepecs, A. Interneuron Types as Attractors and Controllers. Annu Rev Neurosci 43, 1–30 (2020).

22. Asadi, A., Constantinidis, C. & Daliri, M. R. Functional role of cell classes in monkey prefrontal cortex after learning a working memory task. Commun Biol 8, 703 (2025).

23. Pattadkal, J. J., Zemelman, B. V., Fiete, I. & Priebe, N. J. Primate neocortex performs balanced sensory amplification. Neuron 112, 687–688 (2024).

24. Lee, E. K. et al. A multimodal approach for visualizing and identifying electrophysiological cell types in vivo. Nat Commun 17, (2026).

25. Lee, E. K. et al. Non-linear dimensionality reduction on extracellular waveforms reveals cell type diversity in premotor cortex. Elife 10, (2021).

26. Beau, M. et al. A deep learning strategy to identify cell types across species from high-density extracellular recordings. Cell 188, 2218–2234.e22 (2025).

27. Yu, H. et al. In vivo cell-type and brain region classification via multimodal contrastive learning. bioRxiv (2025) doi:10.1101/2024.11.05.622159.

28. Le, T. N. et al. GABAergic Interneuron Differentiation in the Basal Forebrain Is Mediated through Direct Regulation of Glutamic Acid Decarboxylase Isoforms by Homeobox Transcription Factors. J Neurosci 37, 8816–8829 (2017).

29. Anderson, S. A., Eisenstat, D. D., Shi, L. & Rubenstein, J. L. Interneuron migration from basal forebrain to neocortex: dependence on Dlx genes. Science 278, 474–476 (1997).

30. Eisenstat, D. D. et al. DLX-1, DLX-2, and DLX-5 expression define distinct stages of basal forebrain differentiation. J Comp Neurol 414, 217–237 (1999).

31. Potter, G. B. et al. Generation of Cre-transgenic mice using Dlx1/Dlx2 enhancers and their characterization in GABAergic interneurons. Mol Cell Neurosci 40, 167–186 (2009).

32. Zerucha, T. et al. A highly conserved enhancer in the Dlx5/Dlx6 intergenic region is the site of cross-regulatory interactions between Dlx genes in the embryonic forebrain. J Neurosci 20, 709–721 (2000).

33. Soltesz, Z. et al. ACORN: Adaptive Consensus of Repeated Clusterings identifies pyramidal neurons and interneurons in the primate motor cortex based on extracellular spike waveforms. eNeuro 10.1523/ENEURO.0394-25.2026 (2026) doi:10.1523/ENEURO.0394-25.2026.

34. Pachitariu, M., Sridhar, S., Pennington, J. & Stringer, C. Spike sorting with Kilosort4. Nat Methods 21, 914–921 (2024).

35. DeFelipe, J. Types of neurons, synaptic connections and chemical characteristics of cells immunoreactive for calbindin-D28K, parvalbumin and calretinin in the neocortex. J Chem Neuroanat 14, 1–19 (1997).

36. Barbas, H. & Zikopoulos, B. The prefrontal cortex and flexible behavior. Neuroscientist 13, 532–545 (2007).

37. Packer, A. M. & Yuste, R. Dense, unspecific connectivity of neocortical parvalbumin-positive interneurons: a canonical microcircuit for inhibition? J Neurosci 31, 13260–13271 (2011).

38. Barthó, P. et al. Characterization of neocortical principal cells and interneurons by network interactions and extracellular features. J Neurophysiol 92, 600–608 (2004).

39. Kepecs, A. & Fishell, G. Interneuron cell types are fit to function. Nature 505, 318–326 (2014).

40. Golomb, D. et al. Mechanisms of firing patterns in fast-spiking cortical interneurons. PLoS Comput Biol 3, e156 (2007).

41. Hu, H., Gan, J. & Jonas, P. Interneurons. Fast-spiking, parvalbumin^+^ GABAergic interneurons: from cellular design to microcircuit function. Science 345, 1255263 (2014).

42. Markram, H. et al. Reconstruction and Simulation of Neocortical Microcircuitry. Cell 163, 456–492 (2015).

43. Combe, C. L. & Gasparini, S. I from synapses to networks: HCN channel functions and modulation in neurons. Prog Biophys Mol Biol 166, 119–132 (2021).

44. Zaitsev, A. V. et al. Localization of calcium-binding proteins in physiologically and morphologically characterized interneurons of monkey dorsolateral prefrontal cortex. Cereb Cortex 15, 1178–1186 (2005).

45. Helm, J., Akgul, G. & Wollmuth, L. P. Subgroups of parvalbumin-expressing interneurons in layers 2/3 of the visual cortex. J Neurophysiol 109, 1600–1613 (2013).

46. Connors, B. W. & Gutnick, M. J. Intrinsic firing patterns of diverse neocortical neurons. Trends Neurosci 13, 99–104 (1990).

47. Tsodyks, M. V. & Markram, H. The neural code between neocortical pyramidal neurons depends on neurotransmitter release probability. Proc Natl Acad Sci U S A 94, 719–723 (1997).

48. Shaner, N. C., Steinbach, P. A. & Tsien, R. Y. A guide to choosing fluorescent proteins. Nature Methods 2, 905–909 (2005).

49. Ansari, A. M. et al. Cellular GFP Toxicity and Immunogenicity: Potential Confounders in in Vivo Cell Tracking Experiments. Stem Cell Rev Rep 12, 553–559 (2016).

50. Porter, A. G. & Jänicke, R. U. Emerging roles of caspase-3 in apoptosis. Cell Death & Differentiation 6, 99–104 (1999).

51. Duba-Kiss, R., Niibori, Y. & Hampson, D. R. GABAergic Gene Regulatory Elements Used in Adeno-Associated Viral Vectors. Front Neurol 12, 745159 (2021).

52. Rocco, B. R., Sweet, R. A., Lewis, D. A. & Fish, K. N. GABA-Synthesizing Enzymes in Calbindin and Calretinin Neurons in Monkey Prefrontal Cortex. Cereb Cortex 26, 2191–2204 (2016).

53. Kooijmans, R. N., Sierhuis, W., Self, M. W. & Roelfsema, P. R. A Quantitative Comparison of Inhibitory Interneuron Size and Distribution between Mouse and Macaque V1, Using Calcium-Binding Proteins. Cereb Cortex Commun 1, tgaa068 (2020).

54. Brigande, A. M., Krueger, J., Park, C. & Disney, A. A. Calcium-binding protein expression cannot serve as a general classifier for GABAergic neurons in macaque cortex. J Neurosci 10.1523/JNEUROSCI.0192-26.2026 (2026) doi:10.1523/JNEUROSCI.0192-26.2026.

55. Pommer, S. et al. GABAergic interneurons in local circuits of primate prefrontal cortex. Biol Psychiatry 10.1016/j.biopsych.2026.09.012 (2026) doi:10.1016/j.biopsych.2026.09.012.

56. He, J. et al. Machine learning identification of enhancers in the rhesus macaque genome. Neuron 113, 1548–1561.e8 (2025).

57. de Lombares, C., et al. *Dlx5* and *Dlx6* expression in GABAergic neurons controls behavior, metabolism, healthy aging and lifespan. Aging 11, 6638–6656 (2019).

58. Wang, Y. et al. Dlx5 and Dlx6 regulate the development of parvalbumin-expressing cortical interneurons. J Neurosci 30, 5334–5345 (2010).

59. Brown, D. et al. Deep Parallel Characterization of AAV Tropism and AAV-Mediated Transcriptional Changes Single-Cell RNA Sequencing. Front Immunol 12, 730825 (2021).

60. Huang, L.-Y. et al. Characterization of the Adeno-Associated Virus 1 and 6 Sialic Acid Binding Site. J Virol 90, 5219–5230 (2016).

61. Huang, Q. et al. Delivering genes across the blood-brain barrier: LY6A, a novel cellular receptor for AAV-PHP.B capsids. PLoS One 14, e0225206 (2019).

62. Matsuzaki, Y., Fukai, Y., Konno, A. & Hirai, H. Optimal different adeno-associated virus capsid/promoter combinations to target specific cell types in the common marmoset cerebral cortex. Mol Ther Methods Clin Dev 32, 101337 (2024).

63. Aschauer, D. F., Kreuz, S. & Rumpel, S. Analysis of transduction efficiency, tropism and axonal transport of AAV serotypes 1, 2, 5, 6, 8 and 9 in the mouse brain. PLoS One 8, e76310 (2013).

64. Serrano, C., Cananzi, S., Shen, T., Wang, L.-L. & Zhang, C.-L. Simple and highly specific targeting of resident microglia with adeno-associated virus. iScience 27, 110706 (2024).

65. Antinucci, P. et al. A calibrated optogenetic toolbox of stable zebrafish opsin lines. Elife 9, (2020).

66. Senova, S. et al. Experimental assessment of the safety and potential efficacy of high irradiance photostimulation of brain tissues. Sci Rep 7, 43997 (2017).

67. Madisen, L. et al. A toolbox of Cre-dependent optogenetic transgenic mice for light-induced activation and silencing. Nat Neurosci 15, 793–802 (2012).

68. Zerche, M., Wrobel, C., Kusch, K., Moser, T. & Mager, T. Channelrhodopsin fluorescent tag replacement for clinical translation of optogenetic hearing restoration. Mol Ther Methods Clin Dev 29, 202–212 (2023).

69. DeFelipe, J. et al. New insights into the classification and nomenclature of cortical GABAergic interneurons. Nat Rev Neurosci 14, 202–216 (2013).

70. Rudy, B., Fishell, G., Lee, S. & Hjerling-Leffler, J. Three groups of interneurons account for nearly 100% of neocortical GABAergic neurons. Dev Neurobiol 71, 45–61 (2011).

71. Kubota, Y. et al. Selective coexpression of multiple chemical markers defines discrete populations of neocortical GABAergic neurons. Cereb Cortex 21, 1803–1817 (2011).

72. Piatkevich, K. D. & Boyden, E. S. Optogenetic control of neural activity: The biophysics of microbial rhodopsins in neuroscience. Q Rev Biophys 57, e1 (2023).

73. Bansal, H., Gupta, N. & Roy, S. Theoretical analysis of optogenetic spiking with ChRmine, bReaChES and CsChrimson-expressing neurons for retinal prostheses. J Neural Eng 18, (2021).

74. Zhang, C.-J. et al. Effects of fluorescent protein tdTomato on mouse retina. Exp Eye Res 243, 109910 (2024).

75. Taghizadeh, R. R. & Sherley, J. L. CFP and YFP, but not GFP, provide stable fluorescent marking of rat hepatic adult stem cells. J Biomed Biotechnol 2008, 453590 (2008).

76. Medalla, M., Lera, P., Feinberg, M. & Barbas, H. Specificity in inhibitory systems associated with prefrontal pathways to temporal cortex in primates. Cereb Cortex 17 Suppl 1, i136–50 (2007).

77. Medalla, M. & Barbas, H. Synapses with inhibitory neurons differentiate anterior cingulate from dorsolateral prefrontal pathways associated with cognitive control. Neuron 61, 609–620 (2009).

78. Calderazzo, S. M., Busch, S. E., Moore, T. L., Rosene, D. L. & Medalla, M. Distribution and overlap of entorhinal, premotor, and amygdalar connections in the monkey anterior cingulate cortex. J Comp Neurol 529, 885–904 (2021).

79. Mackie, E. C. et al. Mesenchymal cell-derived extracellular vesicles ameliorate age-related deficits in working memory and in vivo MRI measures of white matter structure and function in rhesus monkeys. Geroscience 10.1007/s11357-025-01992-0 (2025) doi:10.1007/s11357-025-01992-0.

80. Siegle, J. H. et al. Open Ephys: an open-source, plugin-based platform for multichannel electrophysiology. J Neural Eng 14, 045003 (2017).

81. Amatrudo, J. M. et al. Influence of Highly Distinctive Structural Properties on the Excitability of Pyramidal Neurons in Monkey Visual and Prefrontal Cortices. J. Neurosci. 32, 13644–13660 (2012).

82. Gilman, J. P., Medalla, M. & Luebke, J. I. Area-Specific Features of Pyramidal Neurons-a Comparative Study in Mouse and Rhesus Monkey. Cereb Cortex 27, 2078–2094 (2017).

83. Ting, J. T. et al. A robust ex vivo experimental platform for molecular-genetic dissection of adult human neocortical cell types and circuits. Sci Rep 8, 8407 (2018).

84. Chang, Y.-M., Rosene, D. L., Killiany, R. J., Mangiamele, L. A. & Luebke, J. I. Increased action potential firing rates of layer 2/3 pyramidal cells in the prefrontal cortex are significantly related to cognitive performance in aged monkeys. Cereb Cortex 15, 409–418 (2005).

85. Luebke, J. I. & Amatrudo, J. M. Age-related increase of sI(AHP) in prefrontal pyramidal cells of monkeys: relationship to cognition. Neurobiol Aging 33, 1085–1095 (2012).

86. Luebke, J. I. et al. Age-related changes to layer 3 pyramidal cells in the rhesus monkey visual cortex. Cereb Cortex 25, 1454–1468 (2015).

87. Medalla, M., Gilman, J. P., Wang, J.-Y. & Luebke, J. I. Strength and Diversity of Inhibitory Signaling Differentiates Primate Anterior Cingulate from Lateral Prefrontal Cortex. J Neurosci 37, 4717–4734 (2017).

88. Aftabjahani, A., Kim, S. J., Wu, C. & Kim, P. Y. High-throughput Image-based Matrix Analysis Platform Software: development of an algorithm for standardized characterization of high-resolution fibrin clot images. Res Pract Thromb Haemost 10, 106795 (2026).

89. Tsolias, A. & Medalla, M. Muscarinic Acetylcholine Receptor Localization on Distinct Excitatory and Inhibitory Neurons Within the ACC and LPFC of the Rhesus Monkey. Front Neural Circuits 15, 795325 (2021).

90. Website. Pavel Kulik (2025). open-ephys-matlab-tools (https://github.com/open-ephys/open-ephys-matlab-tools/releases/tag/v1.3.0), GitHub. Retrieved December 10, 2025.

91. Fortuna, M. G. et al. Histological assessment of optogenetic tools to study fronto-visual and fronto-parietal cortical networks in the rhesus macaque. Sci Rep 10, 11051 (2020).

