## Supplementary Material for "Multimodal characterization of putative *pan*-inhibitory viral constructs in macaque monkeys"

### Supplementary Figures

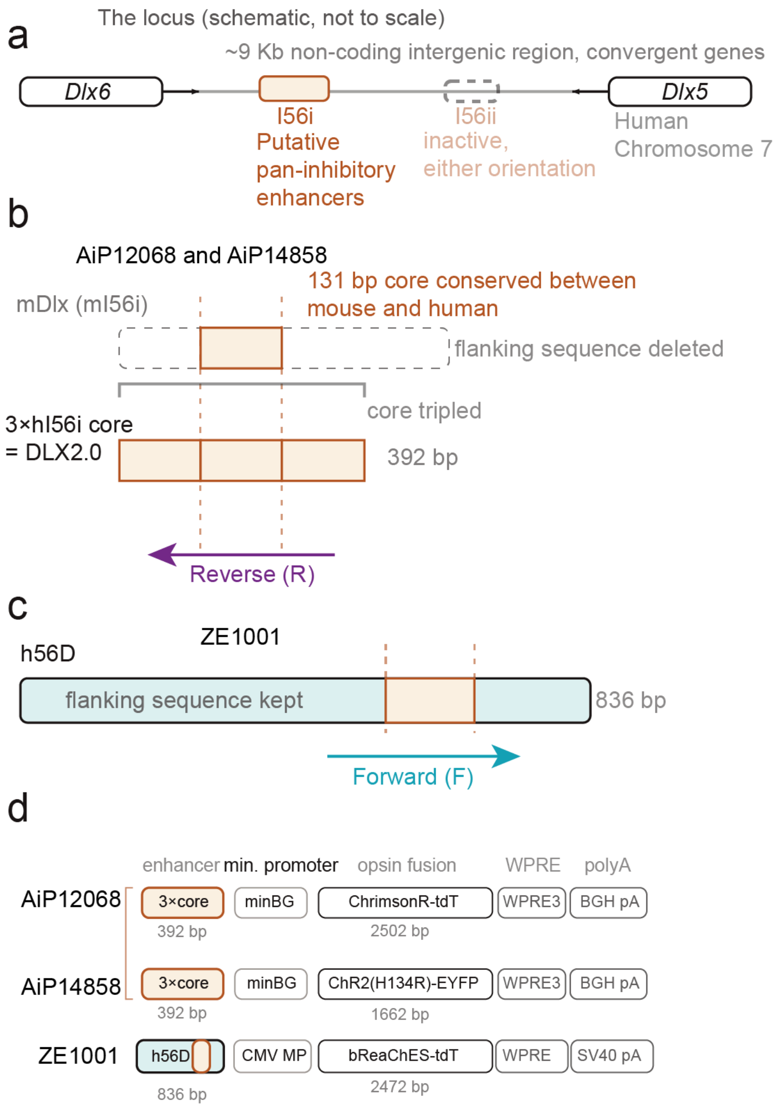


Figure S1: ***Dlx*-based constructs differ in enhancer core copy number, flanking sequence retention, and orientation (a)** Schematic of the human *DLX5–DLX6* locus on chromosome 7, showing the approximately 9-kb intergenic region containing the I56i and I56ii regulatory elements. I56i is a putative pan-inhibitory enhancer; I56ii is indicated as inactive in either orientation. **(b)** Design of the DLX2.0 enhancer used in AiP12068 and AiP14858. A 131-bp I56i core sequence conserved between mouse and human was retained, flanking sequences were removed, and three copies of the human core were combined to generate a 393-bp enhancer in the reverse orientation. **(c)** Design of the h56D enhancer used in ZE1001. The 836-bp sequence retains the I56i core and surrounding flanking sequences in the forward orientation. **(d)** Expression cassette architecture for the three constructs. AiP12068 and AiP14858 combine DLX2.0 with a minimal β-globin promoter (minBG) to drive ChrimsonR–tdTomato or ChR2(H134R)–EYFP, respectively, followed by WPRE3 and a bovine growth hormone polyadenylation signal (BGH pA). ZE1001 combines h56D with a minimal CMV promoter (CMV MP) to drive bReaChES–tdTomato, followed by WPRE and an SV40 polyadenylation signal. Orange boxes denote I56i core sequences; arrows indicate enhancer orientation. Sequence lengths are indicated in base pairs (bp). Schematics are not to scale.

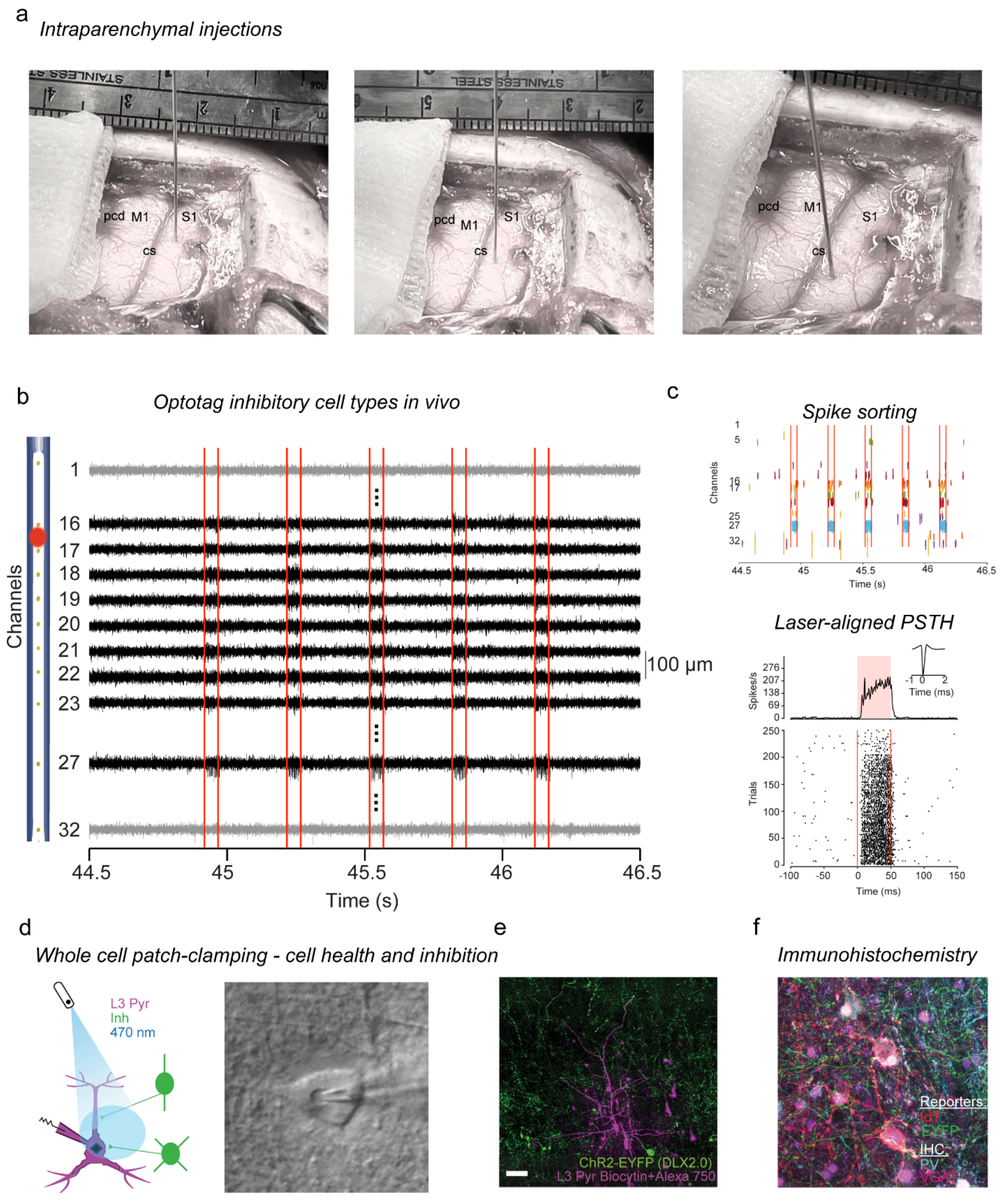


Figure S2: **Reliable extra- and intra-cellular optotagging and optically mediated-coupling pipeline from injection of pan-inhibitory constructs to immunohistochemistry verifies the targeting and activation of subpopulations of inhibitory cells in macaque cortex** (**a**) Three example intraparenchymal injections of the ZE1001 viral construct in Monkey T somatosensory cortex. (**b**) Band-pass filtered data from an anesthetized extracellular recording with monkey B showing a subset of 11 channels from a 32-channel Plexon S-probe with an optical port (located between channels 16/17). Increased neural activity can be observed on channels 16-23 and 27 (emphasized in black) during the stimulation periods (red vertical lines). (**c**) The same time snippet as in **b** demonstrates remaining signal post spike-sorting with Kilosort 4 and manual curation. (*Bottom*) Laser-onset aligned raster plot, trial-averaged peri-stimulus time histogram (PSTH), and average waveform (*inset*) from exemplar optotagged unit recorded from channel 27 in **b** and **c** (*top*). (**d**) Schematic showing *in vitro* whole-cell patch clamp recording in a layer 3 pyramidal neuron (L3 Pyr - purple) synaptically coupled to inhibitory (Inh - green) neurons expressing ChR2 (optically-activated by 470 nm LED - blue). (*Right*) Image of patched pyramidal neuron. (**e**) Intracellular filling with Biocytin of recorded pyramidal neurons. Inhibitory cells expressing Enhanced Yellow Fluorescent Protein (EYFP) are shown in green (seen in the processes of the cells). (**f**) The brain is serially cryosectioned, stained following an immunohistological procedure and four channels are imaged to demonstrate co-labeling of viral construct reporters (e.g., tandem dimer Tomato (tdTomato or tdT)) with specific cell types (e.g., Parvalbumin (PV)) or other markers such as Vesicular GABA Transporter (VGAT).

*Viral construct incubation period (6-20 weeks)*

a

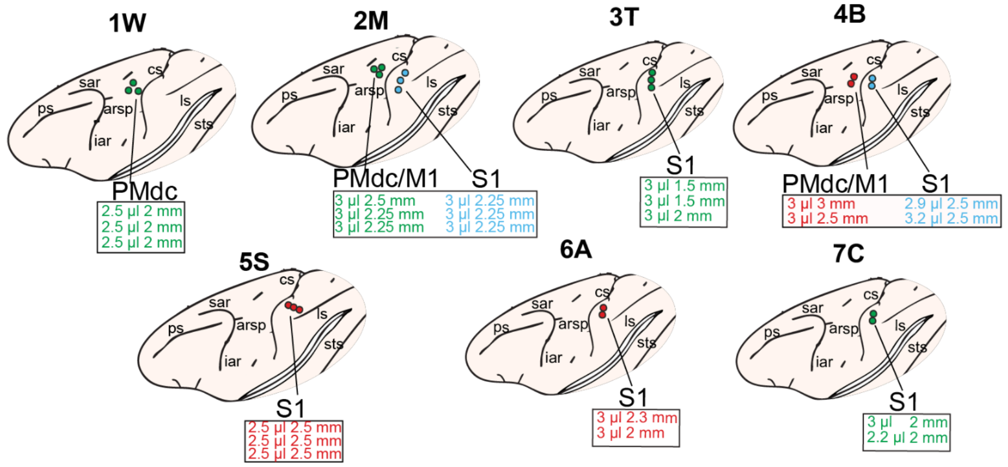


b

*In vivo*

| Construct | ID (Sex Age) | Area | Titer (vg/mL) | Vol. (µL) | Incub. (d) | Opto. | Supp. |
|---|---|---|---|---|---|---|---|
| ZE1001: AAV1/7-h56D-bReaChES-tdT | 1W (F 21y) | L_PMdc | 1 x $10^{14}$ | 7.5 | 96 | 3 | 18 |
| | 2M (M 15y) | L_PMdc/M1 | 1 x $10^{13}$ | 7.5 | 44 | 1 | 2 |
| | 3T (F 16y) | L_S1 | 1.2 x $10^{12}$ | 9 | 63 | 1 | 0 |
| | 7C (F 14y) | R_S1 | 2 x $10^{10}$ | 5.2 | 156 | 13 | 0 |
| AiP1CN4858: AAV/PHP.eB-DLX2.0-ChR2-EYFP | 2M (M 15y) | L_S1 | 7.7 x $10^{13}$ | 9 | 44 | 0 | 0 |
| | 4B (M 13y) | L_S1 | 7.93 x $10^{13}$ | 6.1 | 103 | 11 | 0 |
| AiP12068: AAV/PHP.eB-DLX2.0-ChrimsonR-tdT | 4B (M 13y) | L_PMdc/M1 | 7.7 x $10^{13}$ | 6 | 103 | 16 | 0 |
| | 5S (F 6y) | R_S1 | 1 x $10^{13}$ | 7.5 | 108 | 17 | 0 |
| | 6A (F 16y) | R_S1 | 5 x $10^{12}$ | 6 | 105 | 4 | 0 |

c

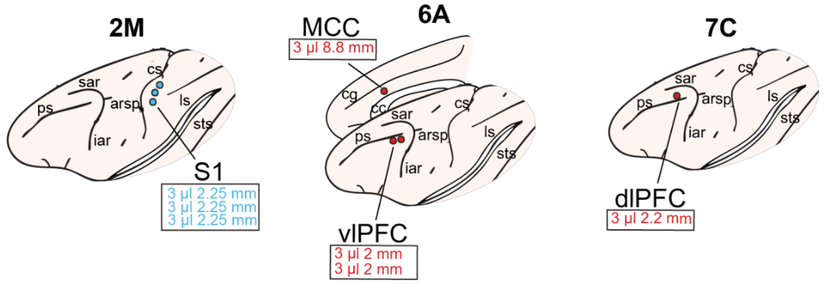


d

*In vitro*

| Construct | ID (Sex Age) | Area | Titer (vg/mL) | Vol. (µL) | Incub. (d) | Opto. | Coup. |
|---|---|---|---|---|---|---|---|
| AiP14858: AAV/PHP.eB-DLX2.0-ChR2-EYFP | 2M (M 15y) | L_S1 | 7.7 x $10^{13}$ | 9 | 44 | 0 | 3 |
| AiP12068: AAV/PHP.eB-DLX2.0-ChrimsonR-tdT | 6A (F 16y) | L_vlPFC | 5 x $10^{12}$ | 6 | 83 | 2 | 5 |
| | 6A (F 16y) | L_MCC | 5 x $10^{12}$ | 3 | 83 | 2 | 4 |
| | 7C (F 14y) | L_dlPFC | 5 x $10^{12}$ | 3 | 93 | 0 | 0 |

Figure S3: **Cortical injection sites and viral construct injection details** (**a**) Schematics (adapted from Fortuna et al, 2020[91]) of 7 monkey brains and the location of viral injections (n=23 in total) illustrated by individual penetrations. Below each schematic brain are the volume (e.g., 2.5 ul) and depth (e.g., 2 mm) of each penetration per color-coded construct. (**b**) Table organized by injected viral construct (Construct) details each monkey's identification (Sex & Age), brain areas injected (Area), titer of viral construct injected (Titer (vg/mL)), total amount of construct injected per area (Vol. (uL)), incubation period in days (Incub. (d)), number of cells optotagged (Opto.) and number of cells suppressed (Supp.). All viral constructs were injected on the same day in a given monkey. If applicable, constructs for *in vivo* versus *in vitro* recordings were injected in opposite hemispheres on separate days (except monkey M, see *4.2*). Constructs can be read here sequentially as viral serotype, enhancer, opsin, and reporter. **c**) Same as in **a** with schematics of 3 monkey brains and the location of viral injections (n=7 in total) separated out by individual penetrations. (**d**) Same as **b** except that the last column details the number of coupled (Coup.) cells. Abbreviations: ps - principal sulcus, sar - superior arcuate sulcus, arsp - arcuate sulcus spur, iar -inferior arcuate sulcus, cs - central sulcus, cg - cingulate sulcus, ls - lateral sulcus, sts - superior temporal sulcus, cc - corpus callosum, cg - cingulate gyrus.

**a**

| Case ID | Area | Viral Construct | R-C (mm) | Volume (mm$^3$) |
|---|---|---|---|---|
| Right hemisphere (*in vivo* recording and serial section IHC) | | | | |
| Mky B | PMdc/M1 | AiP12068 (tdT) | 5.1 | 65.97 |
| Mky S | S1 | | 4.8 | 14.67 |
| Mky A | S1 | | 4.5 | 7.52 |
| Mky C | S1 | ZE1001 (tdT) | 4.5 | 12.67 |
| Mky B | S1 | AiP14858 (EYFP) | 3.48 | 16.04 |
| Left hemisphere (*in vitro* recordings ) | | | | |
| Mky A | MCC | AiP12068 (tdT) | 1.5 | 3.53 |
| | vLPFC | | 1.8 | 2.34 |
| Mky M | S1 | AiP14858 (EYFP) | 3.3 | 11.76 |

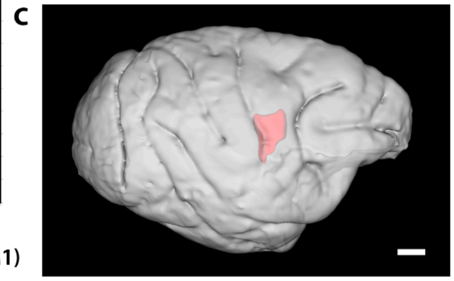


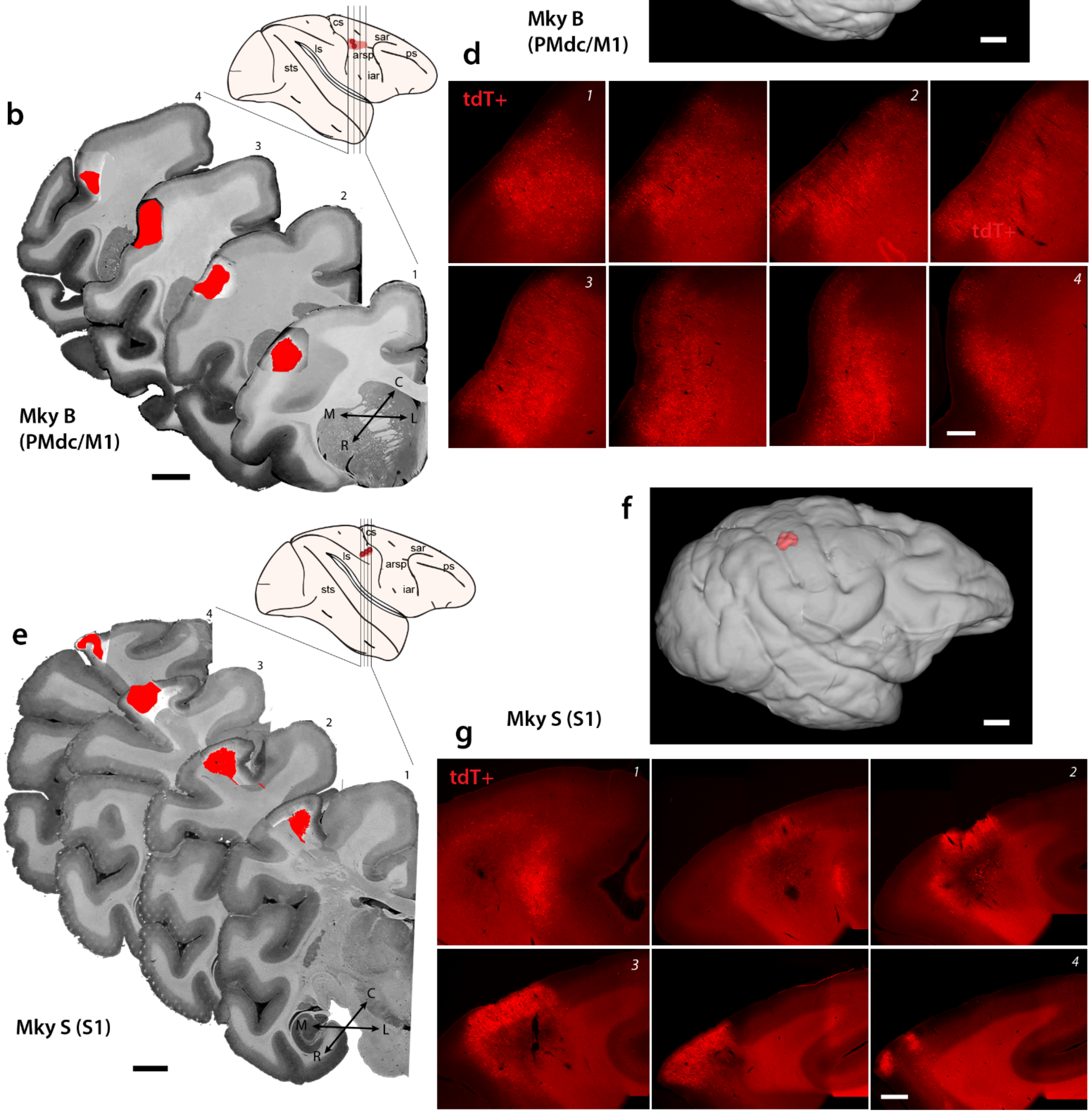


Figure S4: **Injection site reconstruction and validation (a)** Table showing calculated rostro-caudal (R-C) span and volumetric coverage of reporter protein expression within injection sites measured from histological validation and 3D reconstruction analyses. (**b**) Representative serial coronal sections (images 1-4 in **d**) through the rostro-caudal extent of the AiP12068 injection site in PMdc/M1 of Mky B. Lateral brain surface schematic (inset) shows locations of injection site and coronal sections. (**c**) Volumetric 3D reconstruction of the injection site (red) in histological coronal sections registered and projected onto the lateral 3D brain surface based on Mky B's MRI. (**d**) Representative photomicrographs of serial coronal sections through PMdc/M1 of Mky B showing reporter+ protein expression (tdT+)

from the AiP12068 injection site. Images 1, 2, 3, 4 are from rostral to caudal coronal sections shown in **b**. (**e**) Representative serial coronal sections sampled (images 1-4 in **g**) through the rostro-caudal extent of the AiP12068 injection site in primary somatosensory cortex (S1) of Mky S. Lateral brain surface schematic (inset) shows locations of the injection site and coronal sections. (**f**) Volumetric 3D reconstruction of the injection site (red) in histological coronal sections registered and projected onto the lateral 3D brain MRI surface of Mky S. (**g**) Representative photomicrographs of serial coronal sections through S1 of Mky S showing reporter protein expression (tdT+, red) from a AiP12068 injection. Images 1, 2, 3, 4 are from rostral to caudal coronal sections indicated in **e**. Scale bars: **b**, **e**, **c**, **f** = 5mm; **d**, **g** = 1mm.

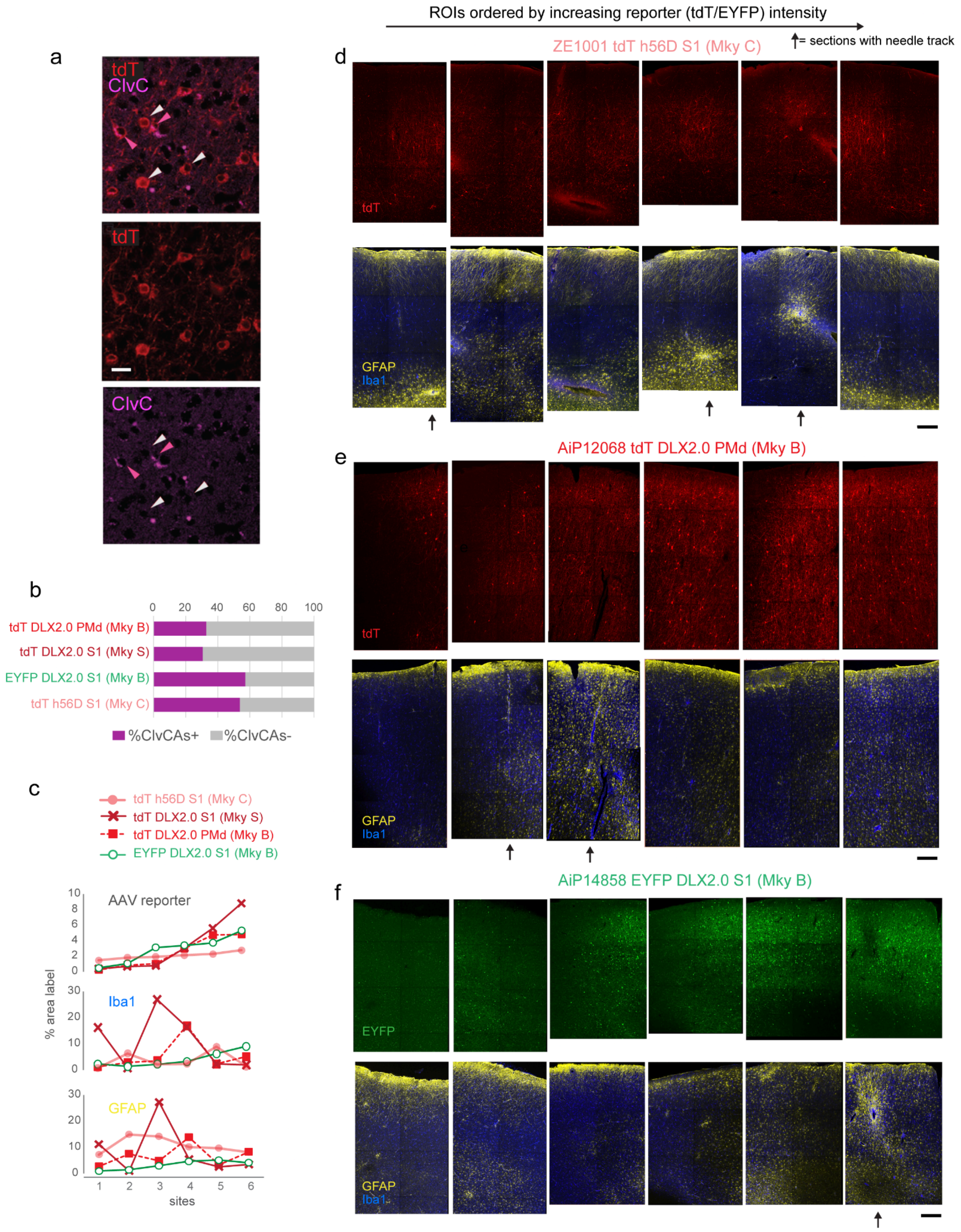


Figure S5: **Majority of construct-labeled cells survive transduction and transgene expression with moderate gliosis present only at the site of construct injection** (**a**) Representative maximum z-projection of a confocal image stack showing tdT+ AiP12068 cells and co-expression of cleaved caspase (ClvC) at high magnification used for quantifying intensity within individual cells. Scale bar = 20 µm. (**b**) Proportion of reporter+ ClvC within each area and case (AiP12068 (tdT DLX2.0): Mky B PMdc/M1, Mky S S1; AiP14858 (EYFP DLX2.0): Mky B S1; ZE1001 (tdT h56D): Mky C S1) (**c**) Percent area label of microglia (Iba1) and astrocytes (GFAP), as markers of gliosis within each ROI site and case. The top graph shows percent area label of reporter protein within each individual ROI site, ordered according to increasing optical density (% area label) of reporter protein expression. *Bottom* two graphs show corresponding glial labeling from each site. Note that density of gliosis markers was not associated with intensity of reporter protein expression. (**d-f)** Representative z maximum-projection of 20x confocal image stack montages of ROIs spanning pia to white matter, within coronal sections sampled across the rostro-caudal extent of the injection site used for analyses in **c**; Top images show single channel tdT (**d**, **e**) and EYFP (**f**) reporter protein label, ordered by increasing intensity of reporter protein. Bottom images show corresponding dual channel GFAP and Iba1 labeling. Arrows indicate sections with the needle track. Scale bars = 200 μm.

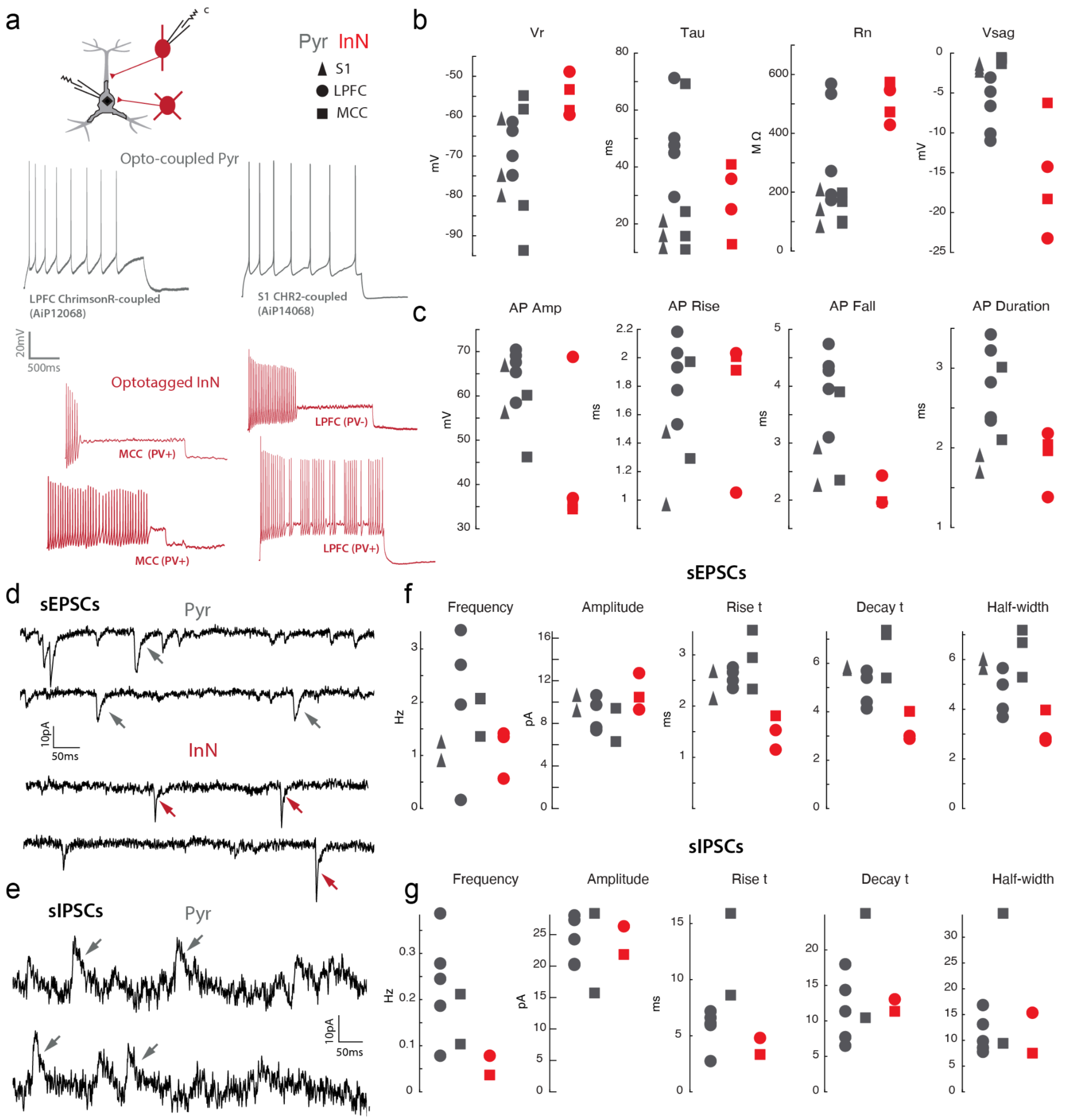


Figure S6: **Biophysical and synaptic current properties of optotagged versus coupled cells** (**a**) *Top*: Schematic of patch clamp recording of a coupled pyramidal neuron (Pyr - gray) and optotagged inhibitory interneuron (InN - red). *Middle*/*Bottom*: Voltage traces showing action potential firing in response to a +100 pA, 2 s current step of coupled pyramidal neurons and optotagged inhibitory interneurons, respectively. Note the regular spiking pattern of pyramidal neurons versus the high frequency and stuttering/phasic firing patterns of inhibitory interneurons. (**b**) Scatter plots showing the resting membrane potential (Vr), membrane time constant (Tau), input resistance (Rn) and sag potential (Vsag) of coupled pyramidal neurons (gray) and optotagged inhibitory interneurons (red). (**c**) Action potential amplitude, rise, fall and duration of coupled pyramidal neurons (gray) and optotagged inhibitory interneurons (red). (**d**) Representative spontaneous excitatory postsynaptic currents (sEPSCs) for pyramidal neurons (*top*) and inhibitory interneurons (*bottom*). (**e**) Two representative spontaneous inhibitory postsynaptic currents (sIPSCs) from coupled pyramidal neurons. (**f, g**) Frequency, Amplitude, Rise time (Rise t), Decay time (Decay t), and Half-width of (**f**) sEPSCs and (**g**) sIPSCs for coupled pyramidal neurons and optotagged inhibitory interneurons.
